# Nucleome Browser: An integrative and multimodal data navigation platform for 4D Nucleome

**DOI:** 10.1101/2022.02.21.481225

**Authors:** Xiaopeng Zhu, Yang Zhang, Yuchuan Wang, Dechao Tian, Andrew S. Belmont, Jason R. Swedlow, Jian Ma

## Abstract

We introduce Nucleome Browser (http://www.nucleome.org), an interactive, multimodal data visualization and exploration platform for 4D Nucleome research. Our tool effectively integrates heterogeneous datasets (e.g., genomics, imaging, 3D genome structure models, and single-cell data) and external data portals by a new adaptive communication mechanism. Nucleome Browser provides a scalable solution for integrating massive amounts of 4D Nucleome data to navigate multiscale nuclear structure and function in a wide range of biological contexts, enabling hypothesis generation and data sharing with the broad community.

## Main Text

The 3D epigenomics community, including the NIH 4D Nucleome (4DN) program [1], has been generating a large number of multimodal datasets with the goal of providing a more complete and multiscale view of nuclear structure and function. Integrating genomic and imaging data as well as analysis results such as 3D genome structure models is of vital importance to creating a comprehensive reference map of nuclear organization. However, effectively exploring these multimodal, heterogeneous 4DN datasets together with other resources for discovery and hypothesis generation is a significant challenge with existing genome browsers. Although specific visualization tools have been developed for different types of nuclear organization data [2–5], no existing tool can effectively integrate genomic, imaging, and 3D genome structure modeling data together with other web-based data portals for scalable and interactive visualization (see **Supplementary Table** S1 and **Supplementary Note** A for a comparison with existing tools).

Here, we introduce the Nucleome Browser (http://www.nucleome.org), an integrative and interactive multimodal data visualization and exploration platform that accelerates access, utilization, and sharing of 4DN data (see **Supplementary Table** S2 for a collection of resources supporting Nucleome Browser, including documentation, tutorial, code repository, etc.). Our platform has two novel capabilities: (1) simultaneous visualization of multiomic datasets (including Hi-C contact maps and 1D genomic signal tracks), imaging datasets, and 3D genome structure models of nuclear organization; and (2) scalable integration of existing data portals in a unified interactive navigation platform, e.g., UCSC Genome Browser [6], WashU Epigenome Browser [7, 8], HiGlass [4], OMERO imaging data server [9], and Image Data Resource (IDR) [10]. A key innovation of Nucleome Browser is an adaptive communication mechanism across multiple web-based visualization web components, which enables users to synchronously explore same data types with linked view, multimodal datasets displayed in web components across different browser tabs, or web applications from different domains. This new design provides a unified interface to explore multimodal datasets and external data portals, and offers a scalable solution for integrating a large number of burgeoning 4DN data and other public genomic and imaging datasets to formulate new hypotheses. Nucleome Browser currently hosts 2,292 genomic tracks and 732 image datasets (see **Supplementary Table** S3 for a summary of 4DN data hosted in Nucleome Browser and **Supplementary Note** B.1 for a tutorial).

Nucleome Browser consists of a series of web components that can be flexibly arranged and customized by the user (i.e., composable and configurable) and that offer synchronized operations for integrative visualization of heterogeneous datasets or different views from the same data modality (**Fig.** 1a, **Methods**, and **Supplementary Note** B.2). Nucleome Browser includes: (1) a genome browser component, designed for visualizing 1D genomic signals (e.g., Repli-seq and TSA-seq [11]) and 2D genomic data (e.g., Hi-C contact maps), (2) a component to display 3D genome structure models (e.g., structure models derived from Hi-C data [12, 13]), and (3) extensible, customized web applications tailored towards specific data types (**Fig.** 1a-b and **Methods** for details). Each web component is self-encapsulated into a panel coupled with the adaptive communication mechanism (**Methods**). As different panels possess the same communication interface, users can flexibly combine and conveniently arrange multiple panels inside one web browser tab for various visualization purposes, including comparisons of multiomic datasets across multiple linked panels (**Supplement Fig.** S23) or multimodal data exploration (**Fig.** 1b and **Methods**). Importantly, the layout of panels can be saved into sessions, enabling management and sharing of the browser’s view from heterogeneous datasets (**Methods**).

**Figure 1.**
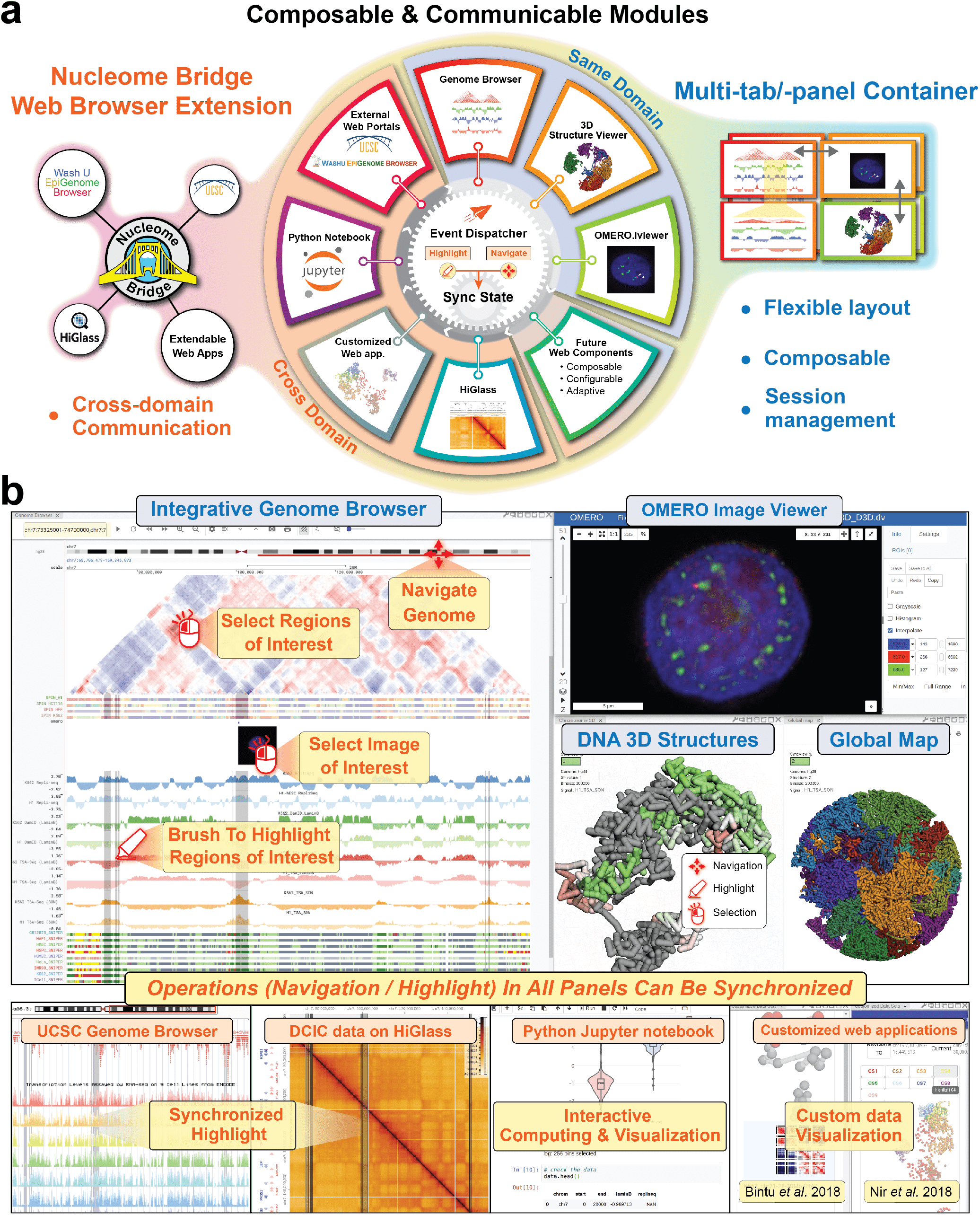
Overall design of Nucleome Browser for integrative and multimodal data navigation. **(a)** Nucleome Browser consists of a series of composable and communicable modules (web components with synchronized operations), each of which is focused on a specific data type. *Middle*: A unified hierarchical event-dispatch system allows modules (outer circle) from either the same domain or different domains to synchronize their status based on genomic coordinates and highlighted regions. *Right*: Web components from the same domain can be combined in a unified panel-container system in a flexible manner supporting synchronization across multitabs. Operations (e.g., navigation to or highlight on the region(s)) initiated in any component will spontaneously dispatch to all connected web components through the event-dispatch system. *Left*: Nucleome Bridge, a web browser extension, empowers various external data portals (e.g., the UCSC Genome Browser) to communicate with other modules in the platform. **(b)** A screenshot of the Nucleome Browser exemplifies interactive exploration of multimodal 4DN data. *Top*: A genome browser web component supports widely-used genomic data types. Users can use the mouse to navigate to or highlight region-of-interest on 1D and 2D tracks. OMERO image viewer shows the details of the image on the selected locus on the image track. 3D genome structure model web component shows a 3D view of chromosome structure with colors indicating the signals of genomic data. The web browser extension named Nucleome Bridge further transmits both genomic coordinates and highlighted regions to other web portals or Jupyter Notebook to generate plots in real-time.

The core component of our adaptive communication design in Nucleome Browser is a multi-channel hierarchical event-dispatch hub (**Supplementary Fig.** S24 and **Methods**), which automatically recognizes a user’s operations triggered in any web component, and then synchronously updates the status of web components or data portals from different domains. For example, when a user navigates to or highlights a region-of-interest in a genome browser panel (**Fig.** 1b), all other panels/portals, including the UCSC Genome Browser, the customized HiGlass viewer, and the Jupyter Notebook, synchronously change to display the corresponding regions, highlight the same region, or update plots in the Jupyter Notebook accordingly (see tutorial in **Supplementary Note** B.3). Note that users can also assign different communication channels to panels to further restrict the synchronization to only happen among panels in the same channel (**Supplementary Fig.** S25). Nucleome Browser is the first platform that supports this multi-domain and multi-channel interactive visualization, thereby greatly enhancing heterogeneous and multimodal data exploration.

To demonstrate the benefits of integrative data exploration using Nucleome Browser, we showcase its unique capabilities with four different data integration examples. **Fig.** 2a shows an example of exploring genome compartmentalization relative to multiple nuclear bodies (i.e., SPIN states [14]) through multimodal data integration (see details in **Supplementary Note** B.4). Specifically, we can select specific SPIN states from a bigBed track and display side-by-side DNA FISH images (from [11]) showing the nuclear localization of these same chromosome regions (**Methods**). The image track consists of the thumbnails of the microscopy images hosted on an OMERO server such that the genomic coordinates of the thumbnails represent the targeted genomic loci of the DNA FISH probes. Users can further explore this FISH data in a pop-out OMERO.iviewer window by clicking a thumbnail (**Fig.** 2a). This example demonstrates the capability of Nucleome Browser for interactive comparison and exploration of genomic and imaging data. Through such integration, users can evaluate the SPIN states based on their proximity to nuclear bodies using DNA FISH image data, thereby connecting genomic readout with what we see from microscopy.

**Figure 2.**
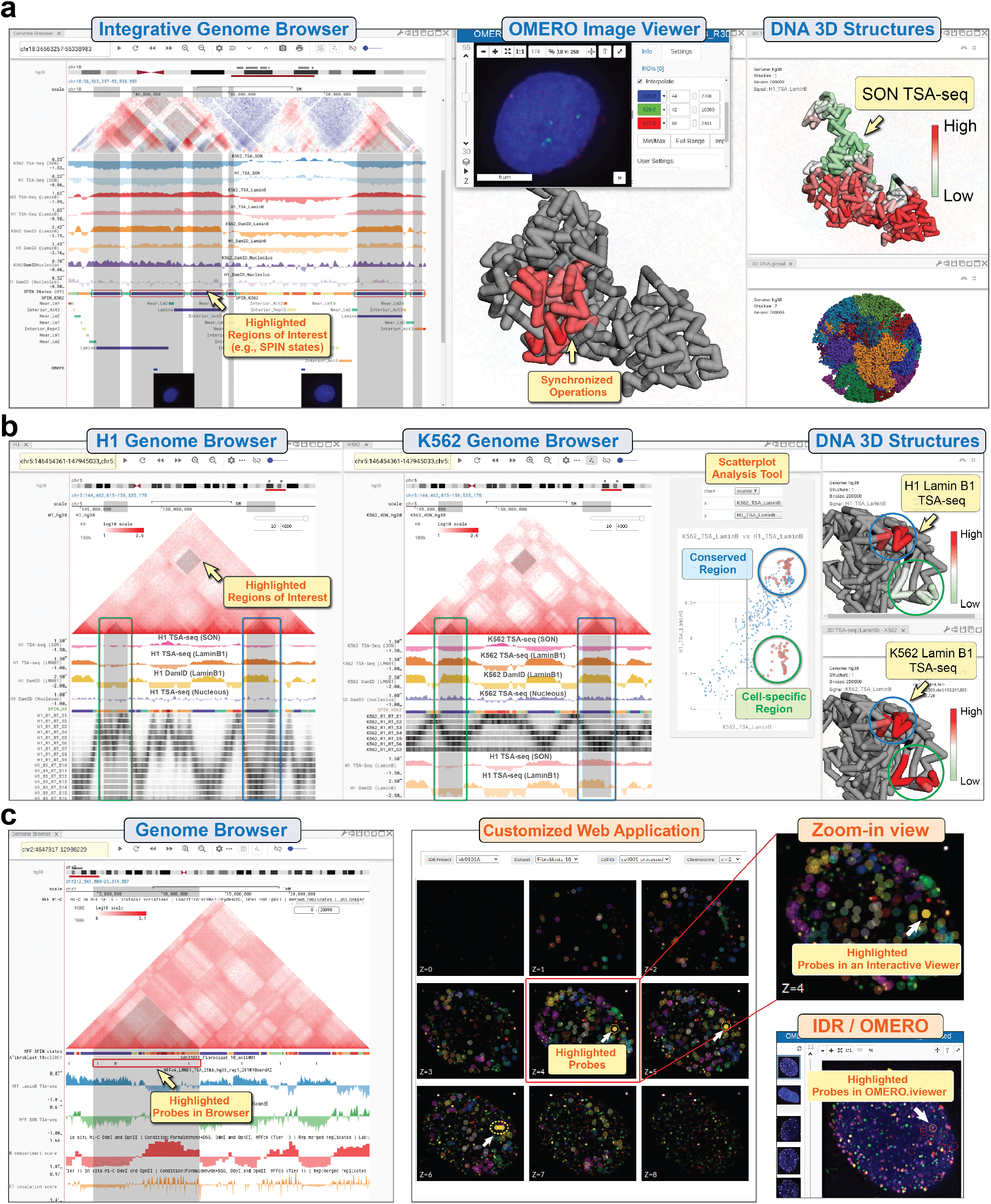
Multimodal datasets and analysis results can be integrated into an interactive, multiscale, and navigable map by Nucleome Browser. **(a)** Nucleome Browser supports integration of genomic and imaging data as well as 3D genome structure models. The genome browser panel on the left shows the input data for the SPIN method [14] in H1 and K562 cells, including TSA-seq and DamID tracks targeting nuclear speckles, nuclear lamina, and nucleolus, bigBed track for the SPIN states, and 2D triangle contact map for Hi-C data. Clicking one category of SPIN states can highlight regions with the same state in the current view. FISH image thumbnails are shown as a track with locations of thumbnail indicate probe loci. Clicking an image thumbnail will visualize the FISH image using the pop-out OMERO.iviewer. 3D genome structure models on the right of the genome browser panel show the overall navigation map (right panel; bottom), highlighted regions of the genome browser panel (middle panel; bottom), and TSA-seq continuous signal (right panel; top), respectively. **(b)** Nucleome Browser facilitates comparisons across multiple conditions. This screenshot shows a comparison between H1 and K562 cells on chromosome 5. Highlighted regions show a cell type-specific LAD/late-replication (K562) to inter-LAD/middle-replicating (H1) transition (green rectangle on the left) and a conserved LAD (blue rectangle on the right) as indicated with TSA-seq, DamID, and high-resolution replication timing Repli-seq tracks. The scatterplot tool and 3D structures further corroborate with the observations from the genome browser panels. **(c)** A customized web application can connect Nucleome Browser to the *in situ* genome sequencing (IGS) data hosted on the IDR platform. Users can navigate the IGS dataset to select individual cell and this web application will automatically fetch images from IDR and mark the location of probes as circles. As users select regions on the genome browser panel on the left, probes overlapping with the highlighted regions will also be highlighted. Clicking a probe will lead users to the IDR portal and open an OMERO.iviewer for further examination.

As a second application, we show how Nucleome Browser supports the synchronized visualization of 3D genome structure models together with genomic and imaging data. **Fig.** 2a displays three different structure model views that integrate features selected in the genome browser panel: (1) Middle panel (bottom): highlighting a chromosome region in the genome browser panel produces highlighting of this same chromosome region in the 3D genome structure model view; (2) Right panel (top): Selecting a SON TSA-seq track in the genome browser panel produces a color-coding display of these SON TSA-seq values superimposed on the 3D genome structure model (SON TSA-seq data from [11]); and (3) Right panel (bottom): a 3D genome global view showing each chromosome in a different color. Importantly, the communication between the genome browser and the 3D structure model panel is bi-directional. Users can select a chromosome segment in the 3D genome model view to directly navigate to this region in the genome browser panel. Alternatively, users can select a chromosome segment in the genome browser panel to visualize this region in the 3D genome structure model view. By integrating these genomic, imaging, and 3D genome structure model views, Nucleome Browser offers an easy, intuitive navigation experience to interactively explore multimodal 4DN datasets. This enables users to generate testable hypotheses of nuclear structure-function connections that may be obscured by individual data modality.

As a third example, we show how Nucleome Browser enables integrative comparisons of nuclear organization in different cellular contexts based on multimodal datasets. In **Fig.** 2b, we compare multiple data types between H1 and K562 cells (**Supplementary Note** B.4), displaying them in side-by-side genome browser panels. As any operation triggered from one panel will be synchronized in the other panels, users can highlight any region-of-interest in one cell type and examine its counterpart in another panel. For instance, in **Fig.** 2b, one highlighted region shows a conserved lamina-associated domain (LAD; blue rectangle) while another highlighted region (green rectangle) shows a cell type-specific LAD/late-replication (K562) to inter-LAD/middle-replicating (H1) transition. We can further add 3D genome structures as an additional evaluation (**Fig.** 2b, right panel). Notably, we have built a scatterplot analysis tool to reveal the global correlation of two bigWig tracks. As shown by the scatterplot in **Fig.** 2b, the cell type-specific region we highlighted is one of several outliers deviating from the generally positive correlation between H1 and K562 Lamin B1 TSA-seq scores. Users can select genomic bins in the scatterplot and highlight them on the genome browser and 3D genome structure models. This same approach to comparative analysis can be applied to single-cell 3D genome data to explore cell-to-cell variation of 3D genome architecture. As proof-of-principle, in **Supplementary Note** B.5 we provide a tutorial showing the procedures to process single-cell 3D genome datasets in [15] and visualize them together with epigenomic data at population level using the Nucleome Browser. **Supplementary Fig.** S22 shows an example of comparing Hi-C contact maps from two single cells along with the predicted 3D genome models.

Recently developed multiplexed immuno-FISH imaging methods [16–20] are now able to provide a direct, whole-genome visualization of spatial 3D chromatin localization that genomic mapping cannot provide. Integrating this multiplexed imaging data with genomic mapping assays and analysis should create a more comprehensive reference map of nuclear architecture. These complex, multimodal data are challenging to present in an intuitive, easy-to-use application. We use our adaptive communication mechanism as the foundation for three new modules that connect Nucleome Browser with external data sources. We built a customized web application to allow users to interactively explore data hosted on Nucleome Browser with *in situ* genome sequencing (IGS) [20] data hosted on IDR (**Fig.** 2c) [10]. Using this web application, users can choose a single-cell image dataset and explore the image data through the OMERO.iviewer together with the markers of probes highlighted on the images. When users highlight chromosome regions in the genome browser, those probes that overlap with the highlighted genomic regions will also be highlighted, allowing users to interactively compare the distance of probes and associated genomic features with their 3D spatial nuclear and chromosomal localization. We established two additional web applications to host single-cell 3D genome structure models derived from multiplexed OligoSTORM [16] and OligoDNA-PAINT [17] data. These web applications allow users to explore the variations among single-cell models that cannot be revealed from population Hi-C contact maps (**Supplementary Fig.** S26a-c). Furthermore, to access public data hosted on the external data portals, we developed a unique web browser extension named Nucleome Bridge to expand data exploration to datasets hosted in the UCSC Genome Browser, the WashU Epigenome Browser, the HiGlass viewer, and a Python Jupyter notebook (**Supplementary Fig.** S26 and **Supplementary Note** B.3). Together, our novel design in supporting integrative exploration across multiple domains has drastically enhanced the capability of visualizing external data sources and can be easily expanded to support new data types.

In summary, Nucleome Browser represents the next-generation design of multimodal data visualization for nuclear organization research. Our tool provides both modular panels for heterogeneous data types as well as user-friendly, multiscale interactive navigation, which will be necessary to meet the growing demands of an upcoming massive wave of new and diverse 3D epigenomics datasets. We anticipate that Nucleome Browser will serve as an important portal for users to fully utilize different types of 4DN and related datasets to formulate hypotheses and provide new insights into the interplay among different components in the nucleus and their roles in nuclear structure and function.

## Methods

### Overall design of Nucleome Browser

Nucleome Browser aims to comprehensively address the challenge of multimodal data visualization for 4D Nucleome research. The key conceptual innovation of multimodal data visualization in Nucleome Browser is that each component has its own purpose but collectively they allow users to achieve integrative and interactive navigation of heterogeneous datasets. In Nucleome Browser, we designed composable (flexible arrangement) and configurable (customized setting) web components, which communicate with each other to achieve synchronized operations, to visualize multimodal datasets (**Supplementary Fig.** S24). Importantly, different types of web components are all self-encapsulated into panels to communicate with each other. For example, a panel (web component) designed for displaying 3D genome structure models could automatically update its view to match the views of other web components. This is achieved by our new multi-channel hierarchical and adaptive communication mechanism (see text below and **Supplementary Fig.** S24), enabling operations triggered by users (e.g., navigation or highlight) in any web component to broadcast to other connected web components across multiple browser tabs (i.e., multi-tabs). In addition, we further developed a web browser extension named Nucleome Bridge as a relay center for real-time cross-domain communication (i.e., multi-domain) among web applications (**Supplementary Table** S1). Notably, our design allows users to synchronously explore multimodal datasets hosted on Nucleome Browser and external data portals/tools, including UCSC Genome Browser [6], WashU Epigenome Browser [7], HiGlass [4], the Jupyter Notebook, and other customized web applications tailored towards specific purposes. The details of different types of web components and adaptive communication mechanism are described below.

### Web components for the visualization of multimodal datasets

Nucleome Browser consists of a series of web components designed for interactive navigation of multimodal datasets, including genomic, imaging, and 3D genome structures as well as customized web applications.

#### Genome browser web component for genomic data

The genome browser web component in Nucleome Browser allows users to explore various genomic and epigenomic data in widely used formats, including bigWig, bigBed, Tabix, and .hic format (**Supplementary Fig.** S24). In the back-end, we implemented a cross-platform tool named nucleserver using GO language to host multiscale genomic data as a web service. Nucleserver utilizes Gonetics (https://github.com/pbenner/gonetics) and bix (https://github.com/brentp/bix) for efficient query of bigWig, bigBed, and Tabix data together with custom scripts to handle data query related to .hic data. In the front-end, our genome browser web component allows users to visualize genomic data through navigation and rendering of the queried data into 1D and 2D genomic tracks. In particular, the genome browser not only supports commonly used operations in existing genomic data visualization tools but also many novel features to facilitate integrative exploration cross multiple panels, including three different visualization modes, an interactive scatterplot analysis tool embedded in the browser, and a drag-and-drop feature to transfer tracks from a genome browser to a 3D genome structure model panel (see **Supplementary Table** S1 for a comparison with existing tools). It also allows users to add multiple web service URLs even for private data with password protection. **Supplementary Note** B.2.1 provides a tutorial and more description on the design of the genome browser web component.

#### 3D genome structure model web component

Integrative visualization of 3D genome structures together with genomic and imaging data would provide users with a three-dimensional viewpoint of the multimodal datasets. We developed a 3D genome structure model web component using the 3Dmol JavaScript library [21] to display 3D genome structures derived from either computational modeling approaches from Hi-C [12, 13]) or recently developed multiplexed imaging methods [18, 20, 22]. This component visualizes 3D genome structures with multiple styles, such as line, cartoon, cross-point, and sphere, providing users an integrative solution to consolidate 3D genome structure models with various genomic data. We created a new data format named nucle3d to store spatial information of 3D genome structure models. The nucle3d data format contains a header section with meta-information (e.g., genome and resolution) and a body section with X, Y, Z coordinates of chromatin segments. Users can use a command-line tool named nucle (https://github.com/nucleome/nucle) to convert 3D genome structure models from other formats such as hss and cmm into the nucle3d format. To facilitate the integrative visualization of multiple 3D genome structure models, we developed several features to allow users to compare 3D models and explore genomic signals in the context of 3D genome structure models. An detailed description of these features and a tutorial can be found in **Supplementary Note** B.2.2.

#### Web applications for a wide variety of customized datasets

In Nucleome Browser, users can create new web components to display customized data and integrate these web components to existing web components using our adaptive communication mechanism (see text below and **Supplementary Fig.** S24). As proof-of-concept, we built four applications to demonstrate the versatility and scalability of Nucleome Browser in integrating new data types and analysis results.

In example I (**Fig.** 2c), we created a customized web application to link Nucleome Browser with *in situ* genome sequencing (IGS) data [20] hosted on IDR [10]. IGS data bridges genomic information of hundreds of probes with their spatial position in single-cell images, which greatly fits the unique capability of Nucleome Browser for integrative visualization of multimodal datasets. We compiled the meta-information of IGS data from human fibroblasts and intact early mouse embryos (IDR project accession #: idr0101) using the IDR API (https://idr.openmicroscopy.org/about/api.html). This web application allows users to select a specific cell, examine *in situ* sequencing images from the IDR platform, and interactively explore the probes’ spatial information on images together with their genomic locations and other data in the genome browser web component. For example, when users navigate in the genome browser, the probes overlapping with the highlighted regions in the genome browser will be synchronously marked on the image.

In example II (**Supplementary Fig.** S26a-b), we obtained the reconstructed 3D genome position data from OligoSTORM [16] from https://github.com/BogdanBintu/ChromatinImaging. This dataset contains the 3D positions of hundreds of 30kb chromatin segments at single-cell resolution across six experiments. Users can select a single cell to visualize its 3D genome structure model and the corresponding 2D distance matrix (**Supplementary Fig.** S26b).

In example III (**Supplementary Fig.** S26c), we downloaded the 3D genome structure models generated by OligoDNA-PAINT [17] from http://sgt.cnag.cat/3dg/datasets. This dataset contains the 3D position of an 8.16 Mb region on chromosome 19 in a diploid genome, separately processed into 9 groups of chromosomal segments (CS1-9) with a resolution of 10kb using the integrative modeling of genomic regions (IMGR) method [17]. Users thus can interactively explore these chromosomal segments together with tracks on the genome browser panel (**Supplementary Fig.** S26a,c).

In example IV (**Supplementary Fig.** S26d), we obtained 3D DNA FISH data corresponding to eight genomic regions (from [11]). We first added these FISH images onto a local OMERO server. We then created a customized image track inside the genome browser web component to display the images as thumbnails based on the genomic locations of FISH probes (**Supplementary Fig.** S27). Users can select any FISH probe to navigate to its genomic location in the genome browser and further click a thumbnail to explore the FISH image data using the OMERO.iviewer interface.

#### Visualizing imaging data in Nucleome Browser

##### (i) Visualizing imaging data from 4DN

We developed a special web component to display imaging data hosted on the 4DN data portal (https://data.4dnucleome.org) [23]. We first compiled meta-information associated with imaging data on the 4DN data portal, extracted imaging datasets containing genomic coordinates information, and saved meta-information into a local database using the binning index algorithm for fast query of images for queries. We then built a customized web component to display imaging datasets as tracks of thumbnails of the microscopy images such that the genomic coordinates of the thumbnails represent the targeted genomic loci of the probes. **Supplementary Fig.** S2 shows a screenshot of viewing DNA FISH datasets generated from [24]. If a user clicks a thumbnail, an OMERO.iviewer window will pop out, allowing detailed exploration of the image data.

##### (ii) Visualizing imaging data from local OMERO server

We also developed a specific plugin of the OMERO server named OMERO-NB-index (https://github.com/nucleome/omero2cnb) to allow users to interactively explore imaging datasets stored on a local OMERO server in a genome browser panel. Once a user adds genomic coordinates meta-information of images as key-value pairs in the OMERO’s PostgreSQL database through its user interface, OMERO-NB-index would monitor the database, store the meta-information into a binning index data structure, and create a web service for efficient query imaging datasets based on a genomic coordinate (**Supplementary Fig.** S27). Users can load this web service in the genome browser web component and navigate image datasets as tracks of thumbnails. When users click an image thumbnail, an OMERO.iviewer will pop out to show the image data.

#### Other web components in Nucleome Browser

**Supplementary Note** B.2 describes the design of other web components and provides a tutorial.

### Multi-channel cross-tab and cross-domain adaptive communication mechanism

#### A panel container system to host heterogeneous panels

We developed a panel container system to host multiple panels based on Golden Layout JavaScript library: https://golden-layout.com. This system has three important features. (1) Flexible layout: Users can open multiple panels and flexibly arrange their layout by resizing the boundary of the panel using mouse or drag-and-drop a panel to set a new layout. (2) Panel management: Users can easily manipulate panels, including duplication, pop-out, rename, closing a panel, etc. We created a panel space widget to save and restore panels saved in local storage. Users can also save the layout of panels into a session and share the session to collaborators using a URL. (3) Cross-panel communication: The panel container system provides a local event-dispatch system, which handles the communication of events across panels in the same domain (as compared with the cross-domain dispatch system supported by the Nucleome Bridge web extension, see below). Notably, this panel container system is flexible and expandable, and can easily incorporate new customized web components to visualize any future data types.

#### A multi-channel hierarchical event-dispatch system with a unified communication interface

We developed a multi-channel hierarchical event-dispatch system that allows different panels or customized web applications to effectively communicate with each other. This design has two components: (1) a unified communication protocol embedded in different panels; and (2) a central event-dispatch hub.

The unified communication protocol defines how different types of panels or web applications handle users’ operations (event emitter) and respond to operations from other panels (event listener). The event emitter and listener from different panels or web applications share a common set of users’ operations (e.g., navigation and highlight), even though they display distinct data types. For example, when users select a region-of-interest in the genome browser panel, the event listener in the 3D genome structure model panel will receive the genomic coordinates and synchronously highlight the corresponding chromatin segments in the 3D genome structure model. Furthermore, both event emitter and event listener have channel ID as their attribute, enabling customized synchronization across linked panels in the same communication channel.

The central event-dispatch hub in our event-dispatch framework adaptively handles communication across panels and web applications (**Supplementary Fig.** S24). It automatically recognizes the source of event emitter and dispatches event message to other panels and web applications in the same communication channel. In the simplest case, this hub uses the event hub interface from the Golden Layout library to support communication across panels on the same web page. Additionally, this design supports communication across web browser tabs hosted under the same domain using the PostMessage HTML5 API and the Broadcast channel API. Lastly, we developed a web browser extension named Nucleome Bridge to enable communication across web applications in different domains. Nucleome Bridge works as a bridge to connect Nucleome Browser with web applications hosted on the allowed URLs once users implement event emitter and event listener in their web applications. This feature provides users the opportunity for integrative visualization of datasets stored in different web-based sources effectively. Note that we can easily expand the allowed URLs to support customized web applications built by users.

Together, this multi-channel, cross-domain, and cross-tab communication system drastically improves the scalability and navigation experience of heterogeneous datasets by defining universal communications across panels, web tabs, and web portals in different domains.

To further allow users to visualize new data types in Nucleome Browser or link other web applications with our platform, we allow users to test their code in CodePen (https://codepen.io), jsfiddle (https://jsfiddle.net), or local server. Once the code is wrapped into a web component, it can be added to the panel system of Nucleome Browser, with a simple configuration change, since all communications across panels or web applications use the same interface. This ease of use and integration is the cornerstone of our multimodal data ecosystem, enabling Nucleome Browser to be highly adaptive to new data types that will emerge in the future.

Note that the communication protocol and the event-dispatch hub have been wrapped into a nb-dispatch JavaScript library (https://github.com/nucleome/nb-dispatch or https://www.npmjs.com/package/@nucleome/nb-dispatch). Nucleome Bridge web browser extension is available at the Chrome Web Store (https://tinyurl.com/nb-bridge).

## Data Acquisition in Nucleome Browser

All datasets available on Nucleome Browser were collected from published work or data portal from public consortium projects. For the public 4DN data, we developed an automatic workflow to download from the 4DN Data Portal (https://data.4dnucleome.org) periodically. We then further curated the meta-information of the data tracks for visualization. In total, Nucleome Browser currently hosts 2,292 genomic datasets from human and mouse and 732 imaging datasets from human and mouse. **Supplementary Table** S4 lists the detailed sources of all datasets used in Nucleome Browser (latest version updated in November 2021).

## Availability of Code, Software, and Documentation

The code of Nucleome Browser is available at: https://github.com/nucleome

Nucleome Browser can be accessed at: http://www.nucleome.org

Documentation and tutorials are available at: https://nb-docs.readthedocs.io/en/latest

Video tutorials are available at: https://tinyurl.com/nb-video-tutorial

## Acknowledgement

Nucleome Browser was first introduced to the community during the 2018 Annual Meeting of the NIH 4D Nucleome Consortium (San Diego, CA). The authors would like to thank members of the 4D Nucleome Consortium and the broad 3D genome community for feedback and suggestions that have improved the tool. The authors are also grateful to members of the Ma lab for helpful discussions and comments on the manuscript. This work was supported by the National Institutes of Health Common Fund 4D Nucleome Program grants U54DK107965 (A.S.B. and J.M.), U54DK107965-05S1 (J.R.S. and J.M.), and UM1HG011593 (A.S.B., J.R.S., and J.M.). J.M. is additionally supported by a Guggenheim Fellowship from the John Simon Guggenheim Memorial Foundation.

## Author Contributions

Conceptualization, X.Z. and J.M.; Methodology, X.Z., Y.Z., and J.M.; Software, X.Z. and Y.Z.; Resources, A.S.B. and J.R.S.; Investigation, X.Z., Y.Z., Y.W., D.T., A.S.B., J.R.S., and J.M.; Writing – Original Draft, Y.Z., X.Z., and J.M.; Writing – Review & Editing, Y.Z., X.Z., A.S.B., J.R.S., and J.M.; Funding Acquisition, A.S.B., J.R.S., and J.M.

