## Supplemental Information for "Nucleome Browser: An integrative and multimodal data navigation platform for 4D Nucleome"

#### A Supplementary Note: Comparison to Existing Tools

Here we describe the conceptual advances and technical innovations in Nucleome Browser. Nucleome Browser aims to solve the daunting challenge of building an efficient platform capable of integrating emerging data types into a single web application to facilitate integrative data analysis, hypothesis generation, and data sharing. We highlight the important advances of Nucleome Browser compared to existing tools.

First, Nucleome Browser achieves efficient visualization of multimodal, heterogeneous datasets within a single platform. For studying nuclear organization, integrating genomic and imaging data together with 3D genome structure models is critically important. However, no existing tool can integrate multimodal datasets for efficient exploration. For example, both the widely used genome browsers (e.g., UCSC Genome Browser [1] and WashU Epigenome Browser [2, 3]) and the platforms specifically designed for visualizing 2D contact maps (e.g., Juicebox [4], Juicbox.js [5], and HiGlass [6]) have limited support to interactively explore 1D and 2D genomic signals together with 3D genome structure models. Notably, even for the same data type, linking and communicating between multiple windows with the same or different zoom factors, which is highly useful to explore genomic loci far away from each other at multiple scales, is not fully supported by existing tools. More recently, multiplexed genome imaging methods [7–11] have shown great potentials to directly measure the spatial position of chromatin in single cells. However, no visualization tool can allow users to view imaging data with results derived from the corresponding genomic mapping assays side-by-side. Nucleome Browser directly addressed such multimodal data visualization challenge in 4D Nucleome research by directly integrating genomic and imaging data as well as 3D genome structures into a unified platform.

Second, Nucleome Browser has a unique adaptive communication mechanism that allows different components to work synergistically for more interactive and integrative visualization. In Nucleome Browser, different data modalities and data sources are represented as distinct web components, with synchronized operations, for efficient data exploration and also for effective development and sharing. Our multi-channel hierarchical communication scheme (in the form of a novel nb-dispatch library) also allows users to create their own web application with the capability to communicate with the rest of the web components in Nucleome Browser. This design is conceptually different from Juicebox and HiGlass that only allow messages transmission across multiple windows with the same type. Specifically, we designed three novel features for synchronization across heterogeneous web components. (1) Both navigation and highlight operations can automatically broadcast across different data modalities (genomics, imaging, and 3D genome structure models). (2) Direct comparison between genomic signals and 3D genome structure models is achieved by superimposing bigWig and bigBed data onto the 3D models with a color-coding display. (3) We created Nucleome Bridge, a web browser extension, to allow Nucleome Browser to communicate with the external data sources or customized web applications, e.g., UCSC Genome Browser and WashU Epigenome Browser. As both data portals host abundant genome annotations, users can use Nucleome Browser to control the navigation on these external web-based tools and explore unique resources provided by them (e.g., the ENCODE data at UCSC Genome Browser and the Roadmap Epigenomics data at WashU Epigenome Browser).

Lastly, Nucleome Browser supports data queries from distributed web locations and achieves secure private data visualization together with public resources. As the amount of data generated from an individual lab grows rapidly, the demand for hosting private servers is common. Several teams have provided their server-less solution to allow users to efficiently visualize data without heavy programming skills [5, 12, 13]. In Nucleome Browser, we developed a cross-platform data service tool to host private data. The only required configuration input is an Excel file or the ID of a publicly-accessible Google Sheet. Once the data service is launched, users can import their private data by adding the address of the data service in Nucleome Browser. Our data service tool can automatically fetch both remote data (e.g., bigBed and bigWig data) stored online and those as local files. Nucleome Browser drastically reduces the learning curve for users to host private data, requiring less computational power on the server end, and more effectively integrating with other datasets.

Overall, Nucleome Browser is a new and unique solution for visualizing multimodal data and also the emerging new data types for 4D Nucleome research.

### B Supplementary Note: Tutorials for using Nucleome Browser

#### B.1 Tutorial I: Exploring 4DN datasets in Nucleome Browser

Here we show how to use Nucleome Browser to explore the 4DN public data hosted on the main Nucleome Browser portal (at Carnegie Mellon University). First, visit the Nucleome Browser by typing http://vis.nucleome.org in the web browser (we recommend using the Chrome web browser for the best experience). You can then choose to open a saved session or a new genome browser panel after you click the "Start" button on the home page.

Once you open a genome browser panel, you can search and add 4DN data tracks in the configuration page by clicking the panel-configuration button on the top-right corner of the panel as shown in **Supplementary Fig.** S1. We have added 4DN public data to the list of default data services; otherwise, you can add it following **Supplementary Fig.** S1. By default, 4DN datasets are grouped by the experiment type and data format. Once you have selected a category of datasets, you can add tracks by clicking the "selection" button. Note that we have manually curated track names to make them easy to understand. You can click the "exclamation button" to toggle a short description of the track. Clicking the "Read more" will lead you to a web page describing this data in the 4DN DCIC data portal. You can also use the search toolbox to search track by the track name. If one experiment has multiple processed data, we add the experiment ID to the track name. You can copy this experiment ID and use the search box to search all tracks associated with this experiment.

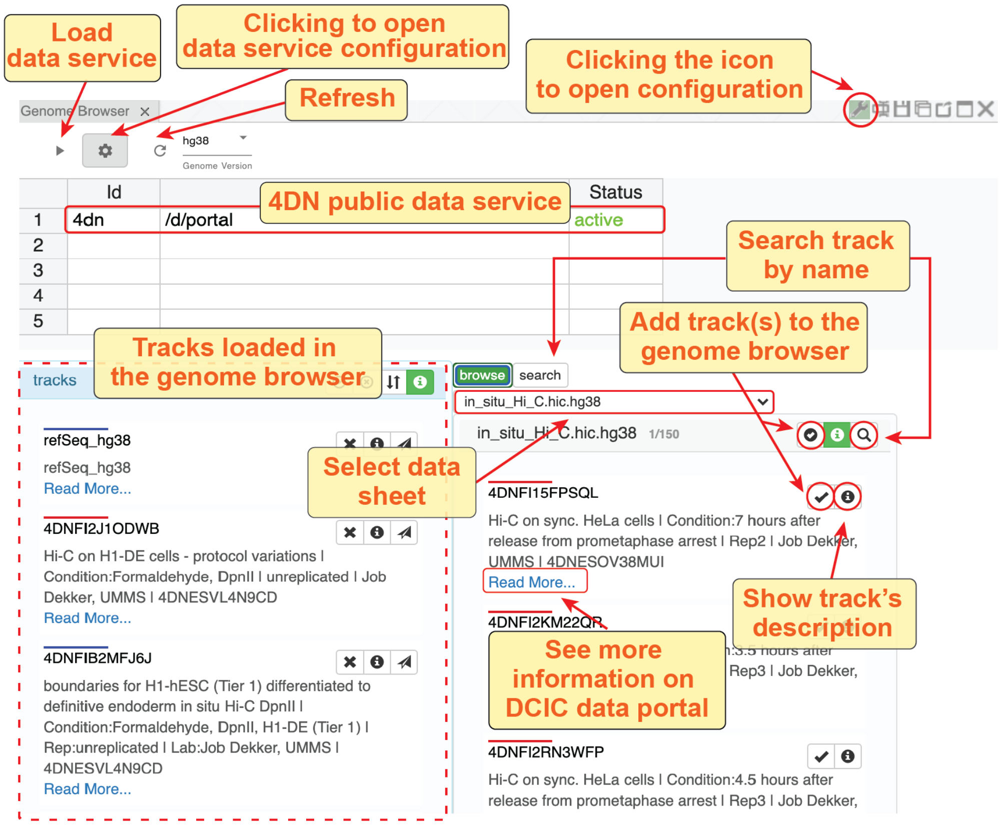


**Figure S1:** The track configuration interface allows users to load public 4DN datasets in the genome browser panel.

Some microscopy imaging data hosted on the OMERO server in the 4DN DCIC data portal are

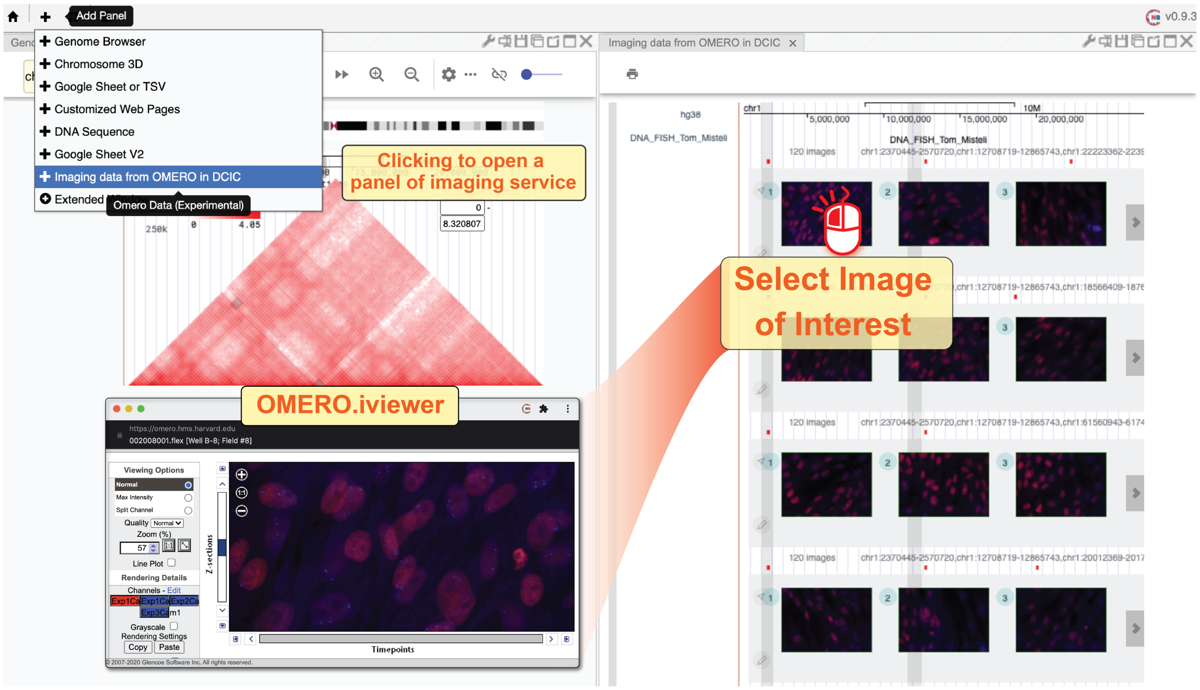


**Figure S2:** Viewing 4DN imaging data using the DCIC imaging data web component in Nucleome Browser.

associated with genomic coordinates information such as the HIPMap data from [14]. To view these datasets, we have compiled a specific web component called "Imaging data from OMERO in DCIC" (**Supplmentary Fig.** S2). In this panel, each track represents one image dataset and individual images from different cells are shown as thumbnails. You can use the arrow keys on each side to browse the thumbnails. The red bars in the track indicate the targeted genomic regions of each experiment from the FISH probes. You can click the thumbnail of an image and you will see a pop-out window leading to an OMERO.iviewer to further examine the image data.

#### B.2 Tutorial II: Novel features of web components to facilitate integrative exploration

In this tutorial, we introduce important features of different types of web components and how to interactively use them for the visualization of mutlimodal datasets.

##### B.2.1 Genome Browser web component for genomic data

We developed several novel features to take full advantage of the composable nature of our web components, which represents significant advances over existing genome browsers.

*(i) Three visualization modes:* In Nucleome Browser, users can open multiple browser panels (web components with the capability to communicate with each other) simultaneously and customize each panel. In particular, we developed three visualization modes to control the synchronization across multiple panels (**Supplemenetary Fig.** S3). (1) Default mode: Both navigation and highlight of region-of-interest synchronize automatically across all genome browser panels. (2) Solo mode: This mode allows users to turn off synchronization in a specific panel, enabling examination of different regions on the same web page. (3) Context mode: This is a variant of the default mode by associating an adjustable zoom factor $K$. In this mode, the current region-of-interest will zoom out $K$ times centered on the region viewed in other panels under the default mode. Therefore, users can explore genomic signals at multiple scales by arranging multiple components side-by-side, one in the default mode and others in the context mode, which is particularly useful for exploring genomic data at region-of-interest and its local context simultaneously.

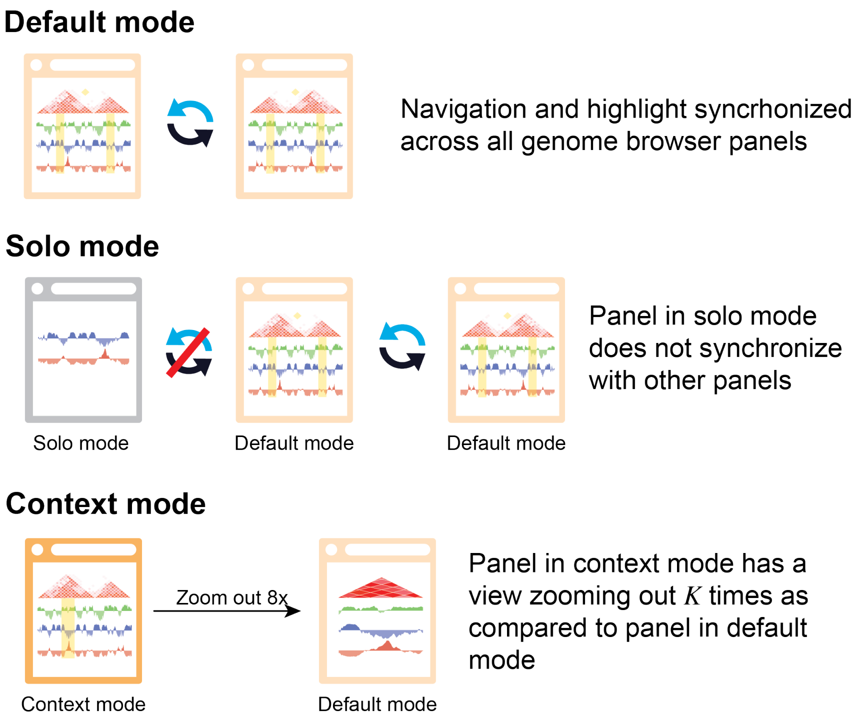


**Figure S3:** Different visualization modes of the genome browser web component. In the default mode, users can use multiple panels, with synchronized explorations, to compare multiomic datasets from different samples. Sole mode allows users to turn off synchronization in one panel, enabling comparison of the same dataset at different genomic loci. Context mode further assigns a zoom factor to each panel such that users can simultaneously explore genomic signals at different scales.

*(ii) Scatterplot analysis tool:* We developed a scatterplot analysis tool in the genome browser web component allowing users to compare genomic tracks quantitatively (**Supplementary Fig.** S4). In the scat-

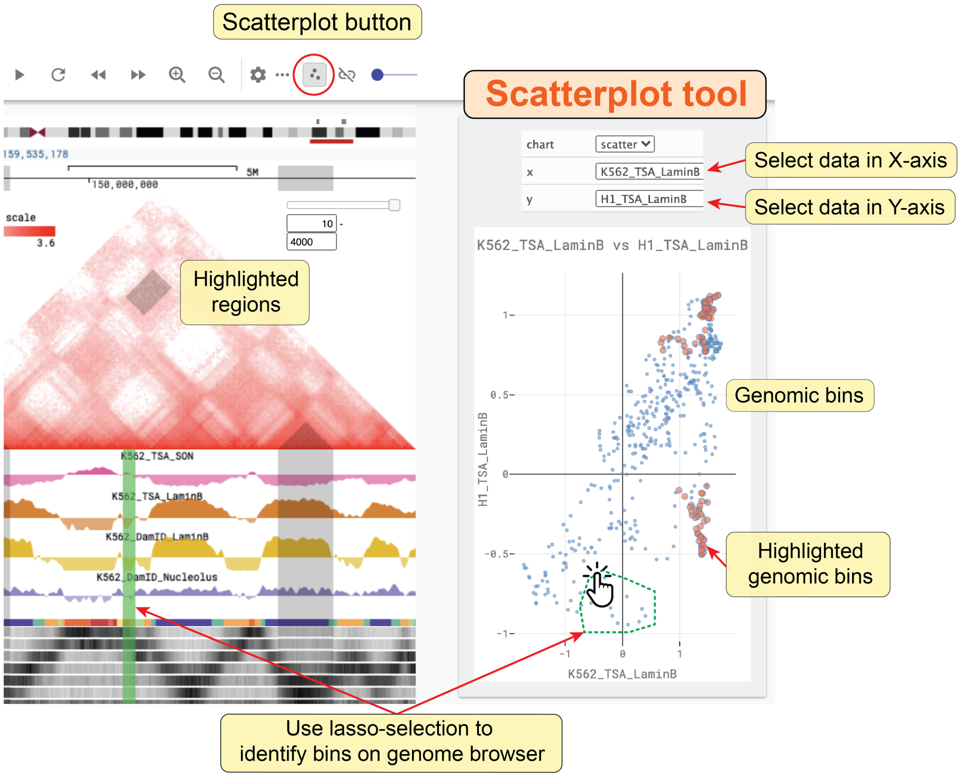


**Figure S4:** Scatterplot analysis tool in genome browser web component. Users can select tracks to show genomic signals on the X-axis and Y-axis of a scatterplot. Highlighted dots (red color) on the scatterplot will be synchronized with the highlighted regions (grey polygon)on the genomic tracks. Users can also use the lasso selection tool to manually pick data points on the scatterplot to view where these genomic bins locate in the genome browser (green polygon).

terplot between two genomic tracks, users can mouse-over dots on the scatterplot to view signal values of two genomic tracks. As users navigate along the genome, the dots would update synchronously. Notably, users can use selection tools to select a group of data points on the scatterplot and the corresponding genomic regions will be highlighted on the genome browser tracks. This operation is reciprocal, i.e., users can also highlight regions on data tracks and view them on the scatterplot.

*(iii) Use drag-and-drop to transfer genomic data to other web components:* We developed a drag-and-drop feature to efficiently share genomic data with other types of web components (**Supplementary Fig.** S5). In the configuration tab of the genome browser panel, users can drag-and-drop a bigBed or bigWig track to a 3D genome structure panel and render the color of 3D genome structure model based on the genomic signals. This new feature offers a way to interactively compare genomic signals with 3D genome structures.

Together, these new capabilities in the genome browser web component in Nucleome Browser can greatly improve user experience of exploring multiomic data tracks.

##### *B.2.2 3D genome structure model web component*

We developed several unique features to achieve effective exploration of 3D genome structure models.

*(i) Modular design:* Similar to the genome browser web component, we encapsulate the 3D genome structure web component into composable and communicable panel, allowing users to open multiple panels simultaneously and visualize the same or different structures with customized settings in each panel.

*(ii) Synchronized view with multi-channel communication mechanism:* Users can assign different com-

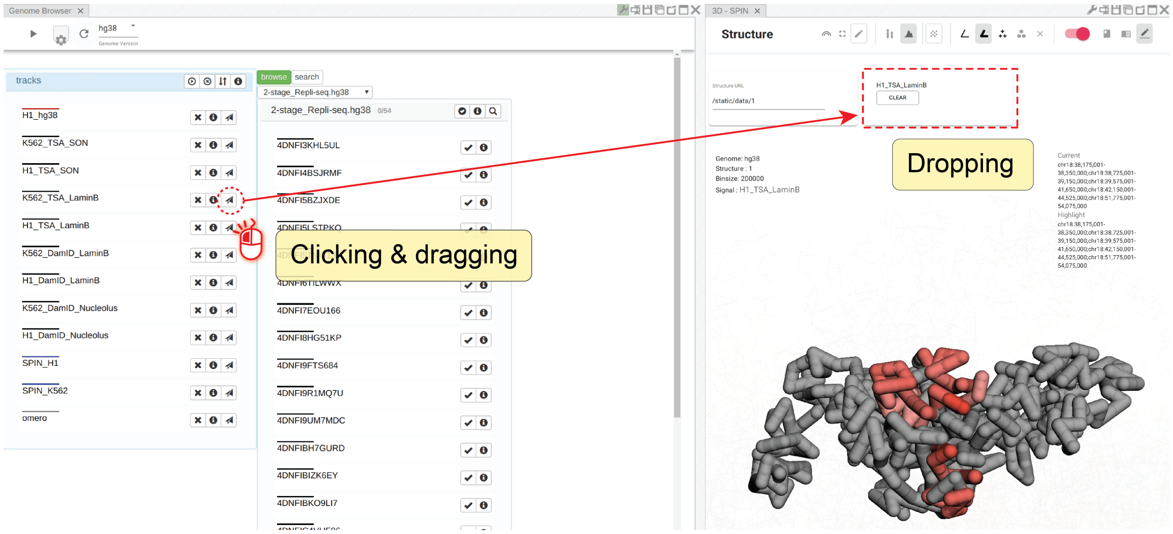


**Figure S5:** Dragging and dropping bigWig track from genome browser panel to 3D genome structure panel to superimpose its values as a color-coding display on the 3D genome structure.

munication channels to each 3D genome structure panel (**Supplementary Fig.** S25). When users visualize the same structure in multiple panels, only structures in panels with the same communication channel ID can synchronize their views including rotation, size, and position of 3D genome structure models. This novel feature is particularly useful when users want to compare different genomic signals superimposed on the same 3D genome structure or examine different spatial contexts of the same genomic region.

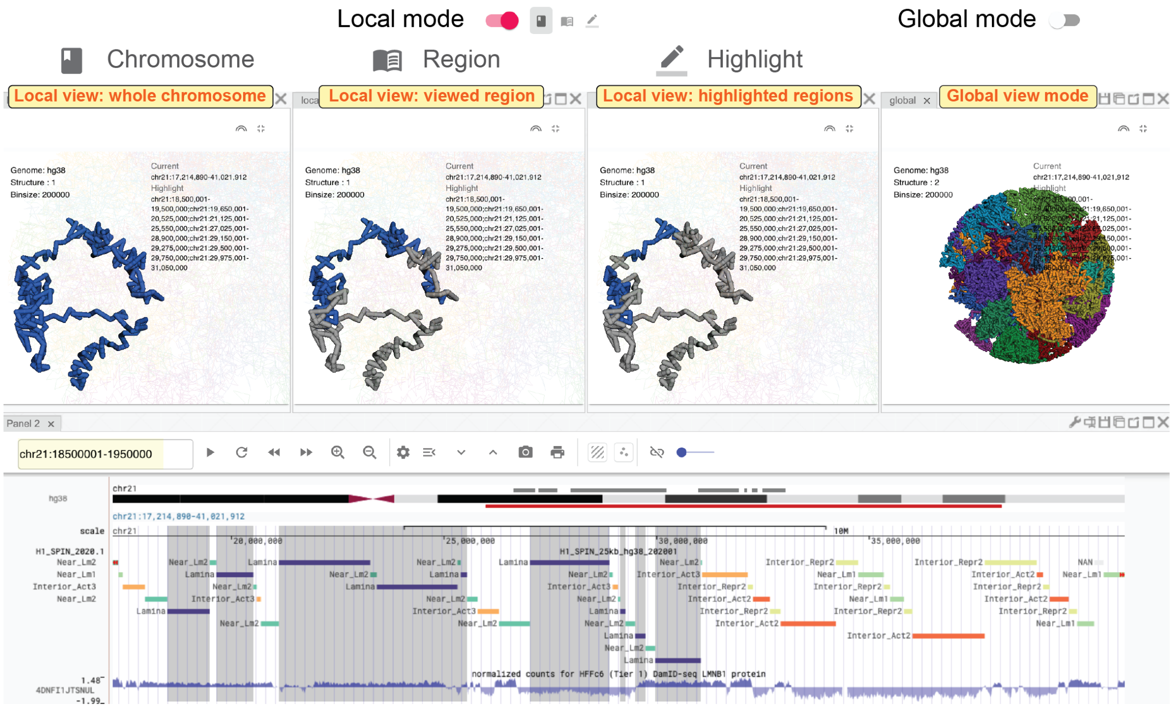


**Figure S6:** In the local model, users can choose different visualization modes to control how 3D models are highlighted relative to the currently viewed region on the genome browser panel. The global mode displays all chromosomes as a global navigable map.

*(iii) Integration with genome browser panel:* The 3D genome structure web component can synchronize its content with information from other web components, e.g., the genome browser web component.

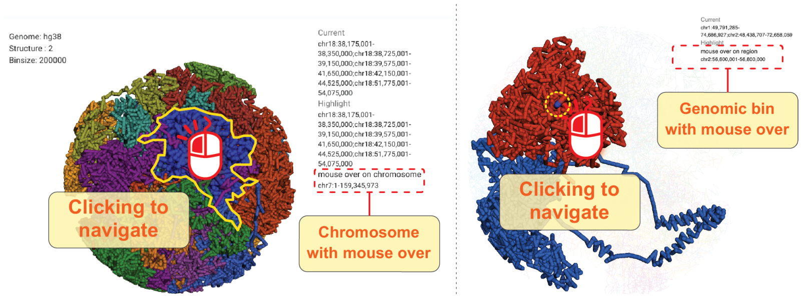


**Figure S7:** Using the global 3D reference panel to navigate the genome. Users can use mouse to select a chromosome to navigate to that chromosome (left). Alternatively, users can use mouse to select a genomic bin to navigate when 3D genome structure model is shown in local mode (right).

Both navigation and highlighted regions are synchronized between the genome browser web component and the 3D genome structure web components. Users can choose three different types of spatial context relative to the currently viewing genomic region, including currently viewing region, highlighted regions, or chromosomes currently being viewed (**Supplementary Fig.** S6).

Notably, a unique design is that the operation is also reciprocal, meaning that users can click segments on a 3D genome structure model to navigate the corresponding genomic regions in the genome browser panel as well (**Supplementary Fig.** S7).

In sum, the 3D genome structure model web component allows users to not only effectively explore multiple 3D genome structures at different scales but also perform integrative visualization with genomic data, imaging data, and customized web applications.

##### *B.2.3 Region-of-interest web component*

The region-of-interest web component works as a convenient tool for users to examine multiple region-of-interest such as ChIP-seq peaks without manually searching for regions (**Supplementary Fig.** S8). Users only need to compile a list of genomic coordinates in a Google Sheet or save them as a tabular text file (tab-delimited) on a web server. This web component then fetches genomic coordinates in the sheet/file, stores them as a local data structure using the binning index algorithm, and renders them as a genomic track. Users can then navigate regions using mouse by clicking each region.

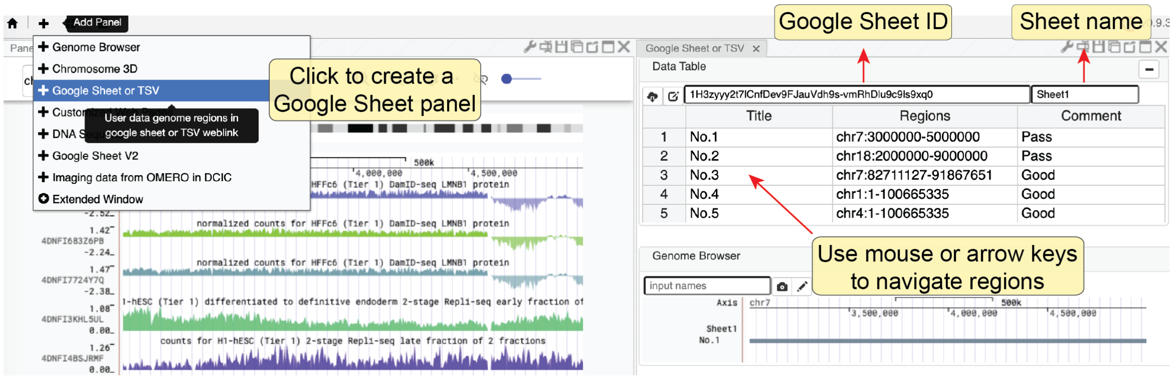


**Figure S8:** Users can link Nucleome Browser with tabular data to efficiently navigate region-of-interest.

#### B.3 Tutorial III: Connecting with external data portals and custom web applications

Nucleome Browser supports not only private data hosted locally by users but it also allows users to use their data hosted on external data portals. Here, we demonstrate how to use the Nucleome Bridge web browser extension to connect Nucleome Browser with UCSC Genome Browser and customized web applications.

##### *B.3.1 Connecting Nucleome Browser with UCSC Genome Browser*

After installation of the Nucleome Bridge web extension in Google Chrome web browser (download link: https://tinyurl.com/nb-bridge), users can click the button of this extension on the web browser to open a synchronized UCSC Genome Browser window. When users highlight some regions or navigate to different regions in the genome browser panel in Nucleome Browser, the UCSC Genome Browser window will jump to or highlight the same regions, accordingly (**Supplementary Fig.** S9). Note that the operation is reciprocal, i.e., the view of the Nucleome Browser will also synchronize with the view on the UCSC genome browser.

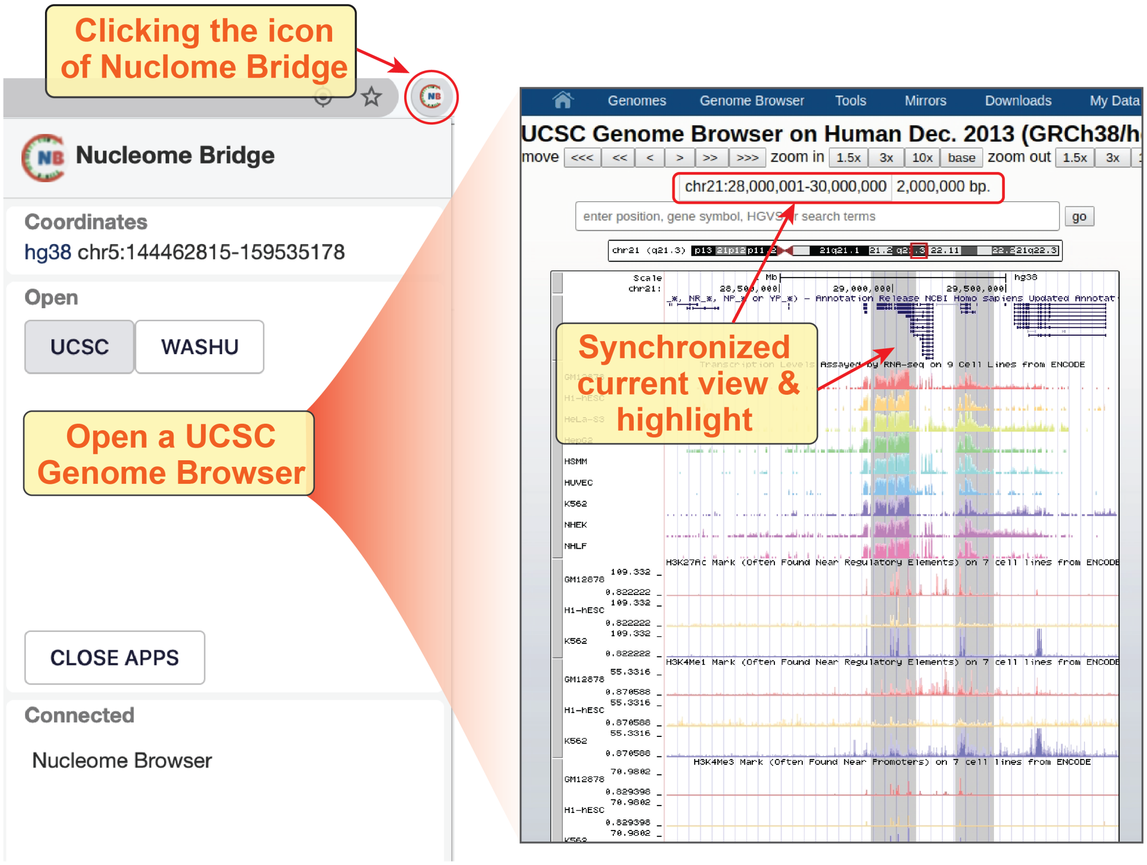


**Figure S9:** Nucleome Bridge web browser extension allows users to connect Nucleome Browser with UCSC Genome Browser, WashU Epigenome Browser, and other web applications.

##### *B.3.2 Connecting Nucleome Browser with Image Data Resource (IDR)*

We built a customized web application to demonstrate that users can interactively explore imaging datasets hosted by IDR together with multimodal datasets in Nucleome Browser as shown in **Fig.** 2c. To interactively explore this example, you can go to https://paper.nucleome.org in a web browser. Click-

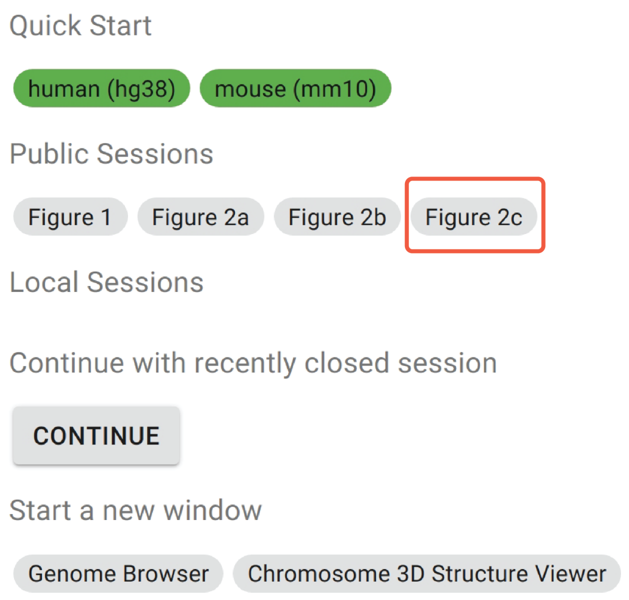


**Figure S10:** Select "Figure 2c" on the quick start page

ing the 'Start' button on the home page, you will see the a quick-start page (**Supplementary Fig.** S10). After clicking the "Figure 2c", you will see a session related to **Fig.** 2c.

**Supplementary Fig.** S11 shows a screenshot of this session. Here, you can select datasets using the dropdown menu in the web application on the left. After you select a dataset, this tool will automatically fetch images and annotation from the IDR website. When you move mouse over a circle (i.e., probe mark), a tooltip will show you the location of this probe and lead you to an OMERO.iviewer if you click the circle. Note that when you highlight regions on the genome browser on the right, probes overlapping with the highlighted regions will be synchronously marked on the image. This application allows users to directly visualize and compare spatial locations of probes and other multmodal data associated with their 1D genomic coordinates.

##### *B.3.3 Connecting Nucleome Browser with HiGlass*

We created a customized web application (http://vis.nucleome.org/static/apps/higlass) to allow users to synchronously view the HiGlass session with Nucleome Browser. This web application allows users to recover the HiGlass session by uploading a configuration file or pasting a session link. Users then can interactively explore datasets on HiGlass together with datasets on Nucleome Browser. For example, when users navigate on the genome browser panel, the HiGlass viewer will jump to the same region (**Supplementary Fig.** S12). Highlighted boxes will synchronously show up on the HiGlass viewer to match the highlighted regions on the Nucleome Browser as well.

We also implemented a drag-and-drop feature to efficiently visualize HiGlass visualization pages on the 4DN DCIC web portal. In any HiGlass display web pages at the 4DN DCIC web portal (e.g., https://data.4dnucleome.org/higlass-view-configs/ae89f1b2-bf71-42db-bee5-e3f7e8b6b590), users can click the 'View JSON' link and drag it on the target box on our web application to directly load the HiGlass session. Our web application will then fetch the HiGlass configuration file automatically from the 4DN DCIC data portal and show the datasets in the HiGlass viewer accordingly.

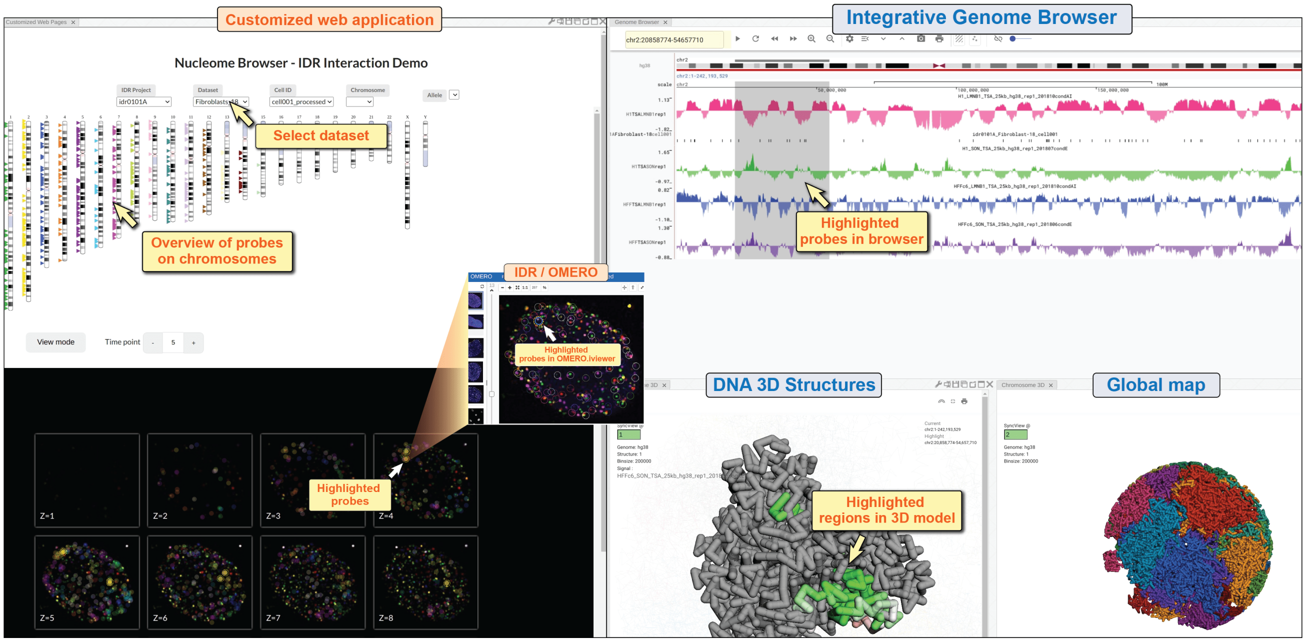


**Figure S11:** Nucleome Bridge web browser extension allows users to connect Nucleome Browser with Image Data Resource (IDR). This screenshot shows a custom web application that allows users to interactively explore *in situ* genome sequencing (IGS) data [11] from IDR together with the multimodal datasets on Nucleome Browser. Video tutorial can be accessed at https://tinyurl.com/nb-video-tutorial.

##### *B.3.4 Connecting Nucleome Browser with Python Jupyter Notebook*

Here, we introduce how to perform interactive data exploration in the Python Jupyter Notebook together with Nucleome Browser. Using the Nucleome Bridge as a relay center, Jupyter Notebook can capture users' operations on Nucleome Browser and update analysis results and plots, accordingly. For example, **Supplementary Fig.** S13 shows that as users change highlighted regions on Nucleome Browser, violin plots update synchronously to display genomic signals under the highlighted regions. Conversely, users can also control which regions to navigate or highlight inside the Jupyter Notebook. This novel feature between the Jupyter Notebook and Nucleome Browser makes data analysis much more interactive and significantly enhances users' productivity. Jupyter notebooks and animations can be found at: https://github.com/nucleome/NB_Jupyter_demo.

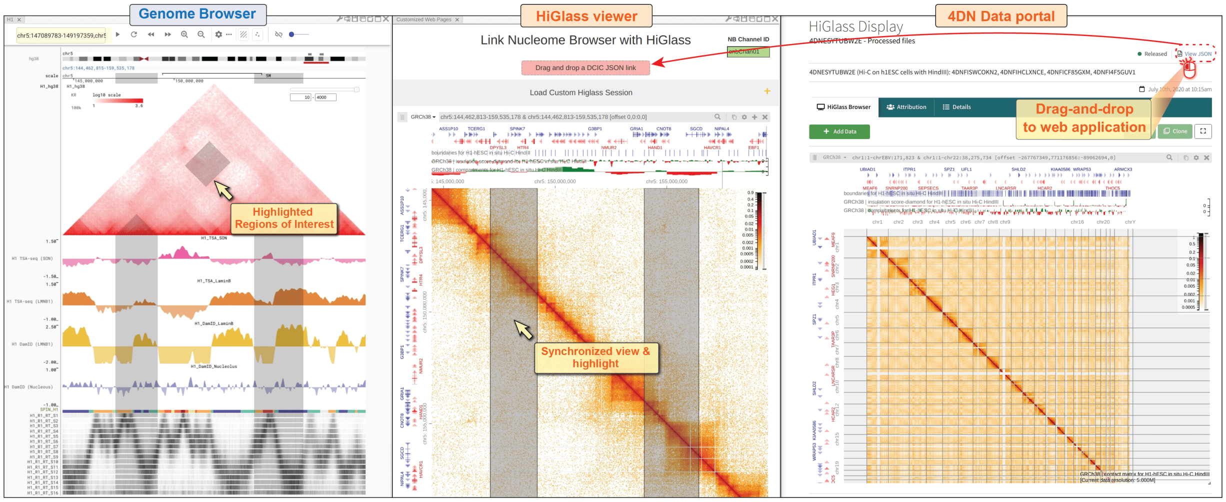


**Figure S12:** Users can interactively explore HiGlass viewer and Nucleome Browser simultaneously. A drag-and-drop feature allows users to efficiently visualize 4DN data on this web application. When users navigate different regions on the genome browser (left), the HiGlass viewer will update to the same region. Highlighted regions are synchronized from Nucleome Browser to the HiGlass viewer. Video tutorial can be accessed at: https://tinyurl.com/nb-video-tutorial.

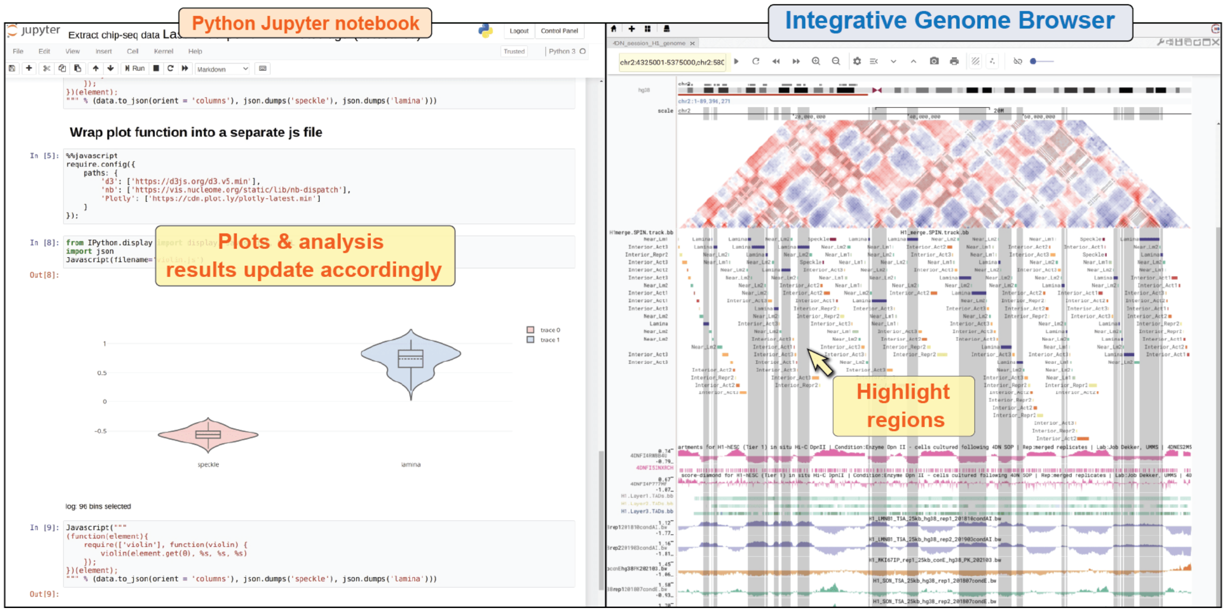


**Figure S13:** Using the Nucleome Bridge web browser extension, users can use Nucleome Browser to interactively perform data analysis on Python Jupyter Notebook. As users highlight different regions on the genome browser on the right, plots in the Jupyter Notebook automatically update to show the distribution of genomic signals based on the highlighted regions. Video tutorial can be accessed at: https://tinyurl.com/nb-video-tutorial.

#### B.4 Tutorial IV: Integrative exploration of multimodal datasets

Nucleome Browser allows efficient and effective integration of different types of data, including genomic, imaging, and 3D genome structure models. Here, we exemplify the advantages of Nucleome Browser over other tools based on sessions related to **Fig.** 2a and **Fig.** 2b in the main text. The goal is to explain various features developed in Nucleome Browser and how we address the challenges of multimodal data visualization.

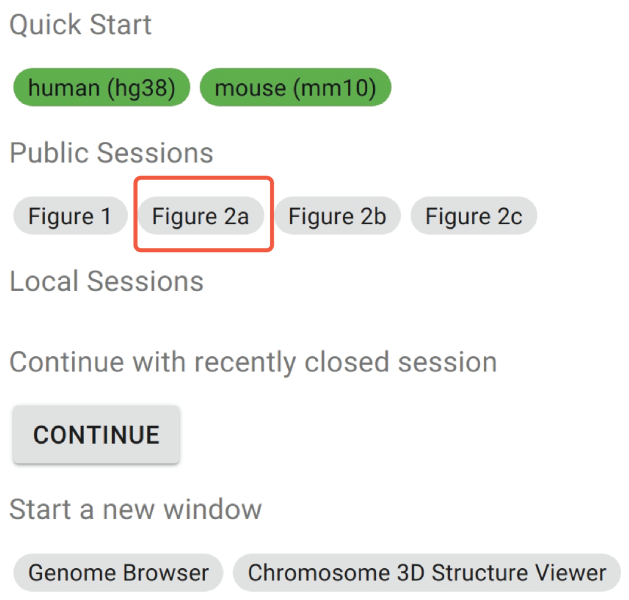


**Figure S14:** Select Figure 2a on the quick start page.

To begin with, you can go to: http://paper.nucleome.org in a web browser to replicate the figures in this manuscript (we suggest using Google Chrome web browser). Clicking the 'Start' button on the home page, you will see the following quick-start page (**Supplementary Fig.** S14). You can click different public sessions related to the figures shown in this paper.

After clicking the "Figure 2a", you will see a session related to **Fig.** 2a showing an example of integrative visualization for different data types (**Supplementary Fig.** S15). The left panel is the genome browser web component showing a series of genomic signals including Hi-C matrix, bigWig, and bigBed tracks as well as thumbnails of images with a blue bar indicating the targeting region of each image. The panel on the bottom right is a global 3D genome reference map showing all the chromosomes in the human genome. The other panels are 3D genome structure panels showing either highlighted regions or the whole chromosome with a color-coding display representing TSA-seq Lamina B1 signals in H1 cells.

Left-clicking on one of the thumbnails of images you will see a pop-out tab showing the details of this image data using the interface of OMERO.iviewer (**Supplementary Fig.** S16).

In the genome browser panel, left-clicking mouse, holding it, and dragging on the genomic tracks, you will see highlighted regions in transparent grey color (**Supplementary Fig.** S17). After you release mouse, you can move the mouse on top of the highlighted region by left-clicking it and holding it. Note that if you highlight an off-diagonal region on the Hi-C contact map, the two associated regions will be both highlighted, accordingly. Additionally, as you move the highlighted regions, the highlighted region on the 3D structure model panel in the middle will also be updated synchronously.

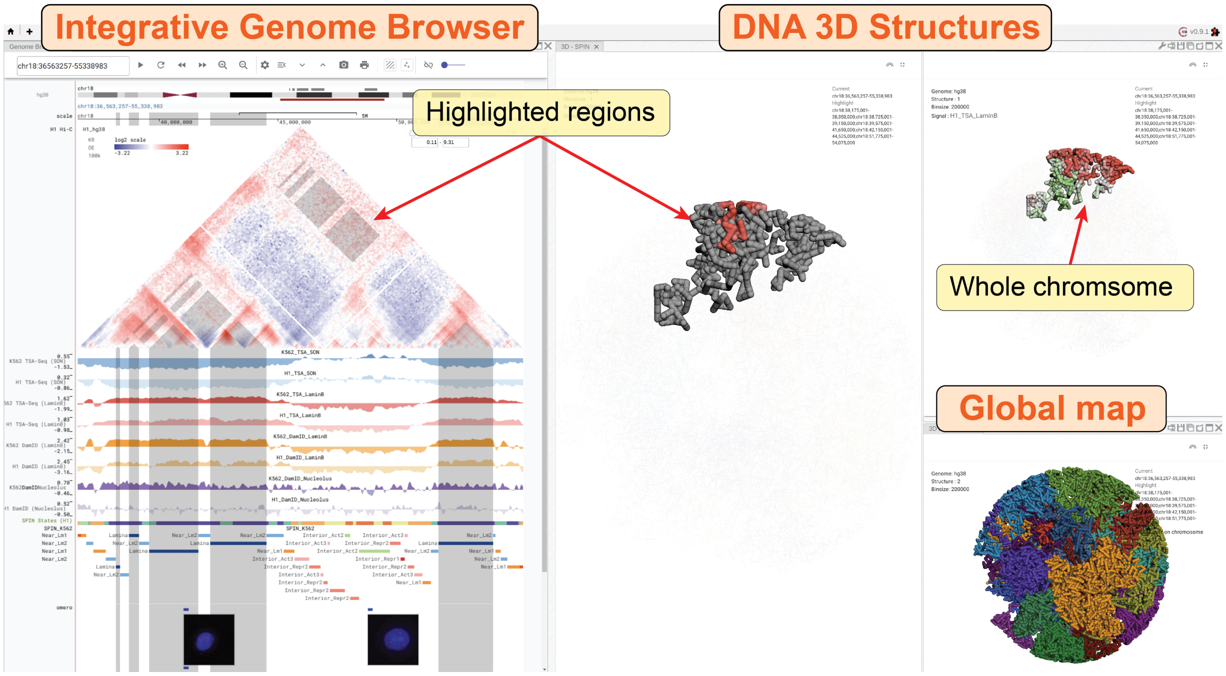


**Figure S15:** An illustration of multimodal data visualization in Nucleome Browser. Video tutorial can be accessed at https://tinyurl.com/nb-video-tutorial.

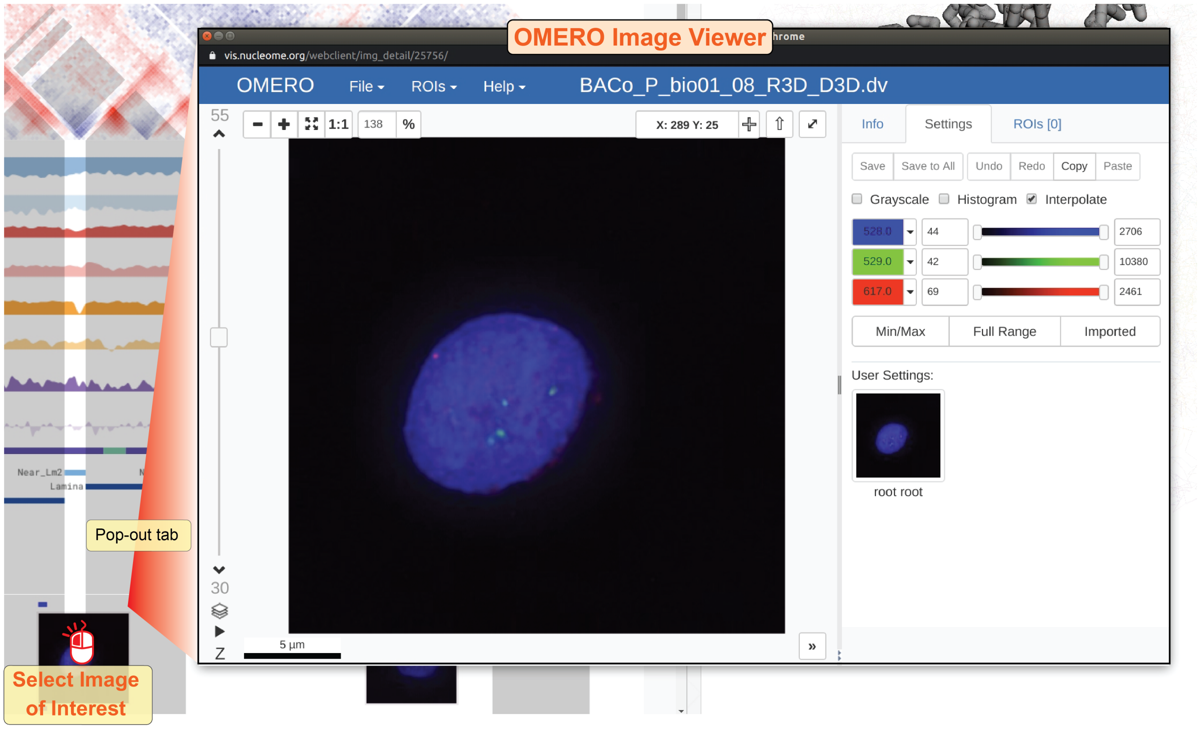


**Figure S16:** An illustration of exploring imaging data in Nucleome Browser by connecting to a local OMERO server.

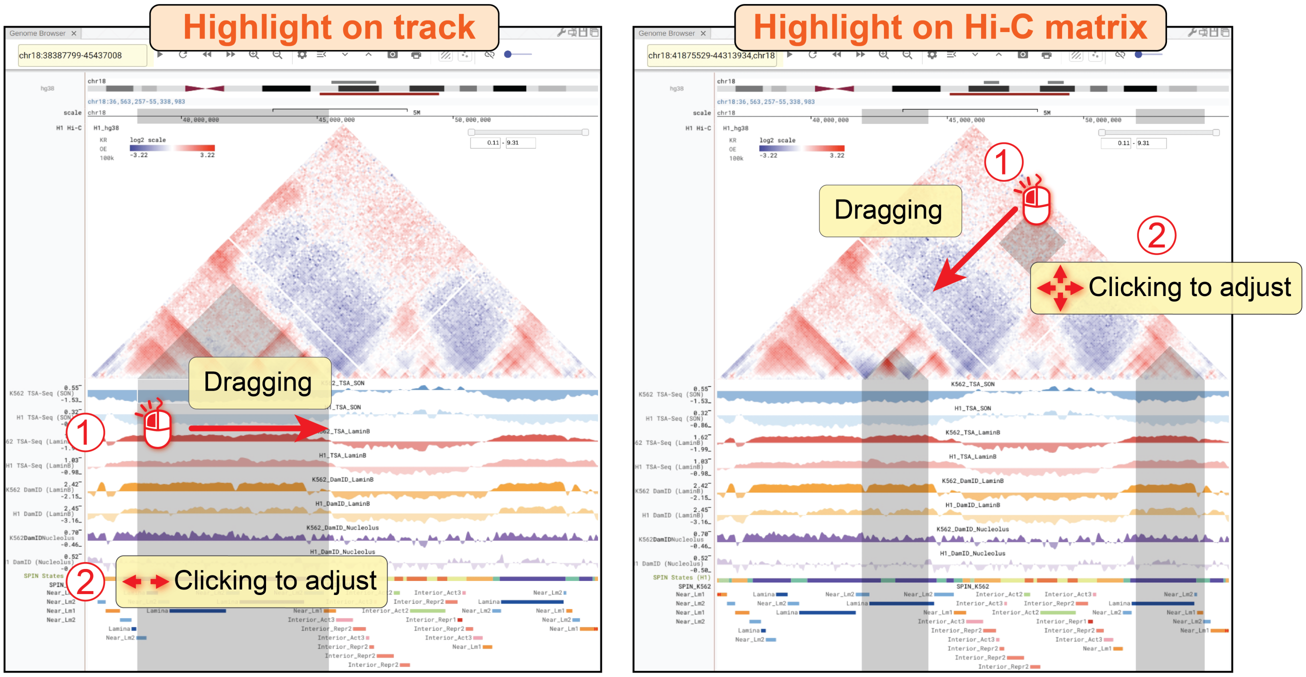


**Figure S17:** Highlighting on genomic data is straightforward. Users can either highlight a single region on 1D genomic signal track or two genomic loci on 2D track such as Hi-C contact map. Users can click mouse on the highlighted region (grey polygon) and drag to further adjust it.

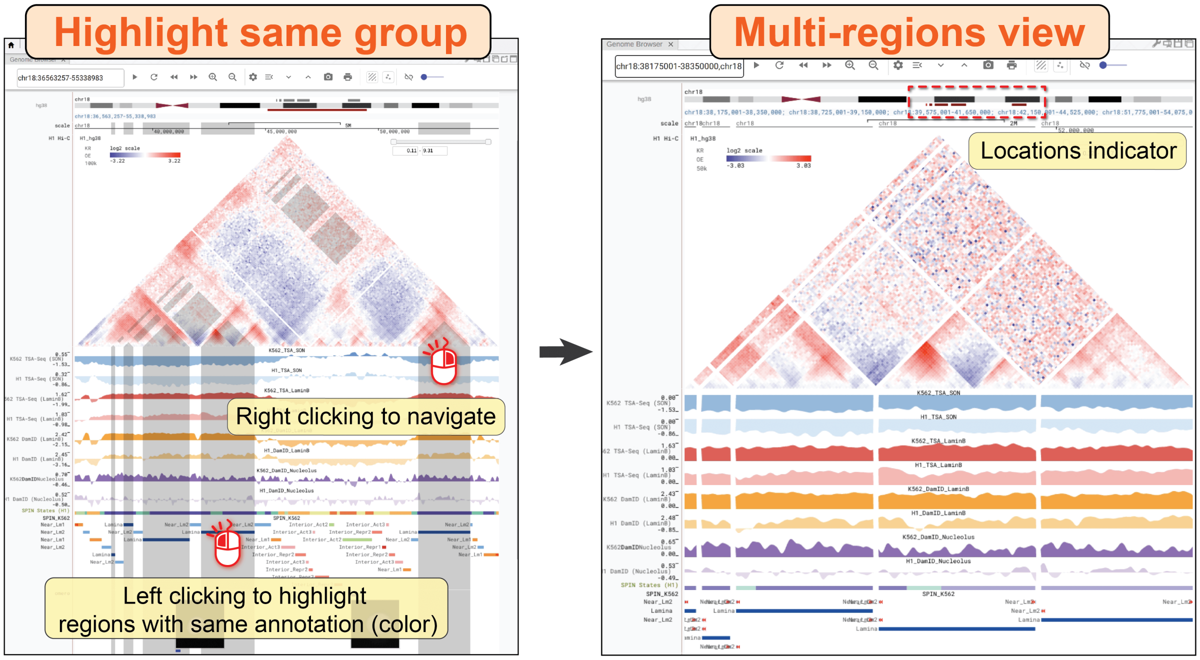


**Figure S18:** Clicking a region on the bed track, other regions with the same color will be highlighted simultaneously. This feature is useful to visualize annotations of chromatin segmentation such as SPIN [15]. When right-clicking the highlighted regions, you will navigate to the highlighted regions (in multi-regions view) even though they are not consecutive.

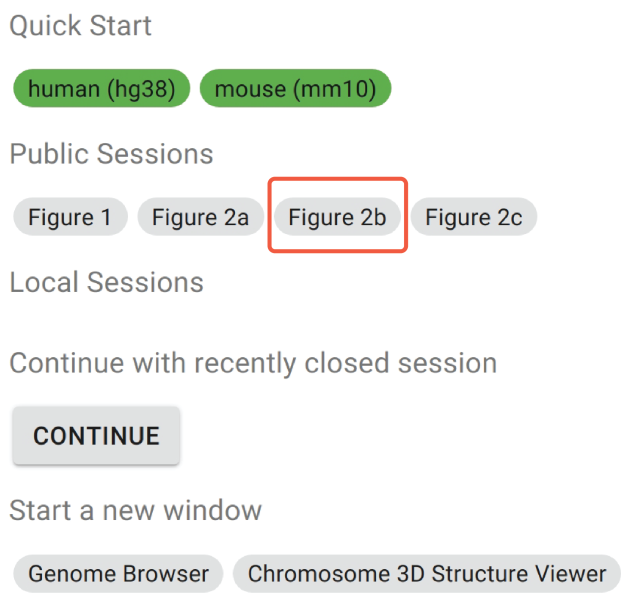


**Figure S19:** Select "Figure 2b" on the quick start page

You can click a region shown in bed tracks to highlight all the regions with the same color (**Supplementary Fig.** S18). Here, the colors can be defined with the 9th column of the bed file following the definition of 12-column bed format: https://genome.ucsc.edu/FAQ/FAQformat.html#format1.

Upon right-clicking the highlighted regions, you will be directed to the highlighted region(s). If highlighted regions are not consecutive on the genome browser you will see stitched regions connecting multiple 'sliced' highlighted regions (similar to the multi-region used in the UCSC Genome Browser) (**Supplementary Fig.** S18).

Comparison of 3D genome structures and their roles in nuclear function between different cell types or conditions is highly informative. Nucleome Browser is an ideal tool to achieve this type of comparison because of its modular architecture. For example, users can create multiple panels with each of them representing one condition or cell type. Users then can view or highlight regions in a synchronized manner across multiple panels. Here, we illustrate how to effectively compare multimodal data between two cell types using the session related to **Fig.** 2b. In the quick-start page on the home page, after clicking the 'Figure 2b' button, you will see a session associated with **Fig.** 2b showing a comparison of various genomic data between H1 and K562 cells (**Supplementary Fig.** S19).

In this session, two genome browser panels on the left show a comparison of various genomic data between K562 and H1, including Hi-C contact matrices, TSA-seq, and DamID tracks mapping to multiple nuclear bodies (e.g., nuclear speckles, nuclear lamina, and nucleolus), and DNA replication timing profiles (**Supplementary Fig.** S20). Two 3D genome structures on the right connect 3D structures with genomic signals by rendering the chromatin according to TSA-seq Lamina B1 signal in H1 and K562, respectively. Importantly, you can highlight any regions on the genomic tracks and compare them to other cell types. For example, the default highlighted regions in this session indicate two dramatically different states: 1) a region showing a conserved lamina-associated profile in H1 and K562 as marked by the blue rectangle; 2) a region showing a cell type-specific pattern between H1 and K562 as marked by the green rectangle. In H1, the cell type-specific region has a relatively lower lamina B1 TSA-seq signal than K562, which is consistent with its earlier replication timing profiles compared with the late replicated profile in K562.

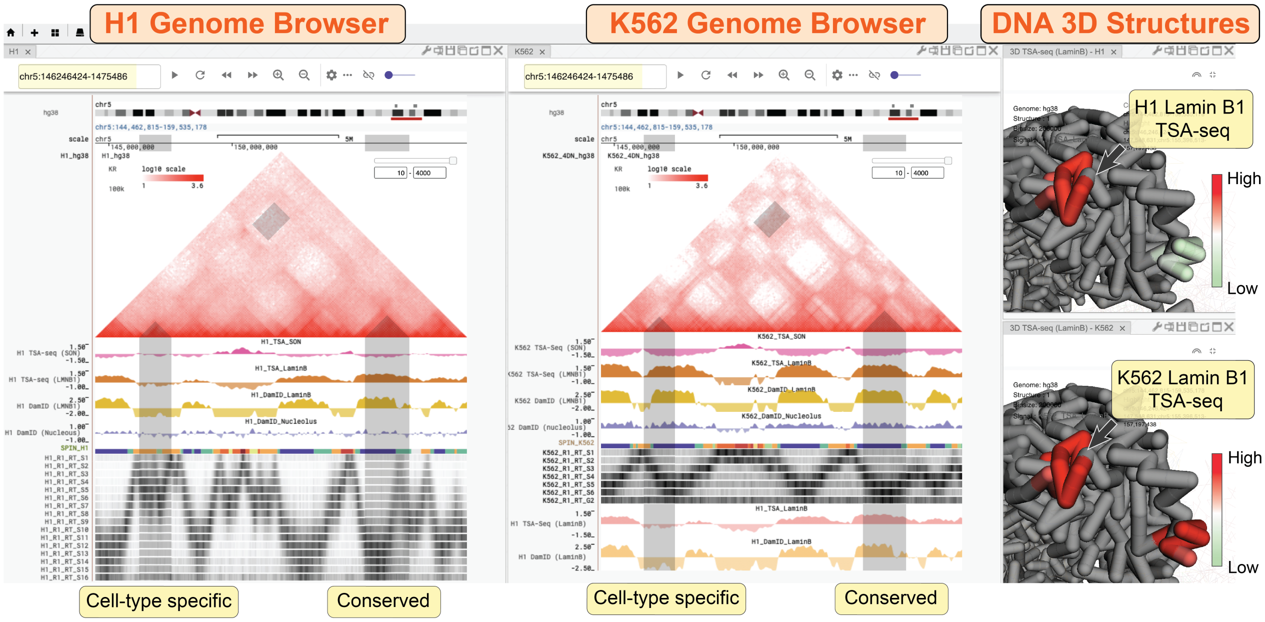


**Figure S20:** Nucleome Browser facilitates the comparison of nuclear structure-function connection between different conditions and cell types. Video tutorial can be accessed at: https://tinyurl.com/nb-video-tutorial.

Left-clicking any place in the 3D genome structure panel, holding the mouse, and dragging it you will see different views of the 3D genome structure (**Supplementary Fig.** S21). You can use the scroll wheel of the mouse to zoom-in and zoom-out the 3D structure. Holding the control key and left-clicking and holding the mouse you can move the 3D structure inside the panel.

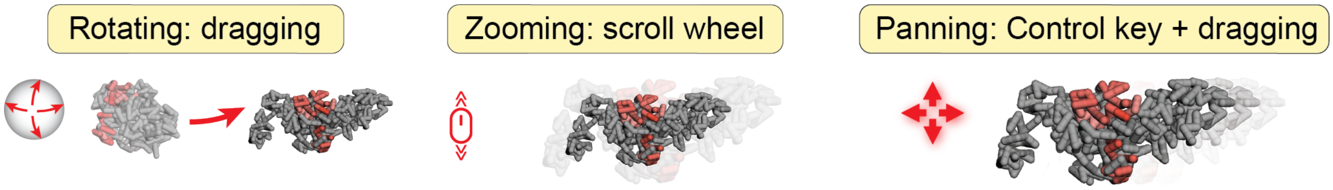


**Figure S21:** Users can pan, move, and zoom-in/out on the 3D genome structures.

##### B.5 Tutorial V: Building a custom data service - An example using single-cell Hi-C data

Here we show how to set up a custom Nucleome Browser session by launching a local genomic service using published data (population Hi-C and single-cell Hi-C) [16] downloaded from gene expression omnibus (GEO) under the accession number GSE80280. In [16], the authors generated population Hi-C data from the mouse embryonic stem cells, eight single-cell Hi-C datasets from the same cell type as well as 3D structure models inferred from Hi-C using the Chrom3D tool [17]. The whole script used to process the downloaded data can be accessed at: https://github.com/nucleome/Tutorial-SingeCellHiC.

###### *Getting data from GEO*

We first create a new folder and use wget command to download the whole dataset from GEO to a local computer and named it 'GSE80280_RAW.tar' as shown below:

```
# create new a folder
mkdir single_cell_hic
cd single_cell_hic
# Download data
wget -O GSE80280_RAW.tar "https://www.ncbi.nlm.nih.gov/geo/download/?acc=GSE80280&
    format=file"
```

Next, we extract files from the tar file in the current directory:

```
# Extract file
tar -xvf GSE80280_RAW.tar
gzip -d *.gz
```

You should see multiple files in different formats with various suffixes, such as .bed, .pdb, etc.

###### *Pre-processing of ChIP-seq peaks*

In order to launch data service and visualize these data in Nucleome Browser, we need to pre-process data and convert them into standard formats supported by Nucleome Browser. First, we use tools from the UCSC Genome Browser and Blat application binaries built (http://hgdownload.cse.ucsc.edu/admin/exe/) to convert ChIP-seq peak files into bigBed format as shown below:

```
# Fetch chrom.sizes information
fetchChromSizes mm10 > mm10.chrom.sizes

# Add additional optional BED fields
# Replace "ChIP-Seq" with sample name
awk '{print $1"\t"$2"\t"$3"\t"$4"\t0\t.\t"$2"\t"$3"\t255,0,0"}' ChIP-Seq_peaks.bed
    > ChIP-Seq_peaks.detail.bed

# Create bigBed file
bedToBigBed ChIP-Seq_peaks.detail.bed mm10.chrom.sizes ChIP-Seq_peaks.bb
```

###### *Pre-processing of population Hi-C and single-cell Hi-C data*

The Hi-C contact map must be transformed into .hic format in order to display in Nucleome Browser. Here we use Juicer tools (https://github.com/aidenlab/juicer) to create .hic files from the downloaded contact pair files. The Hi-C contact pair file should be converted into one of the formats as shown in the documentation of Juicer tools (https://github.com/aidenlab/juicer/wiki/Pr).

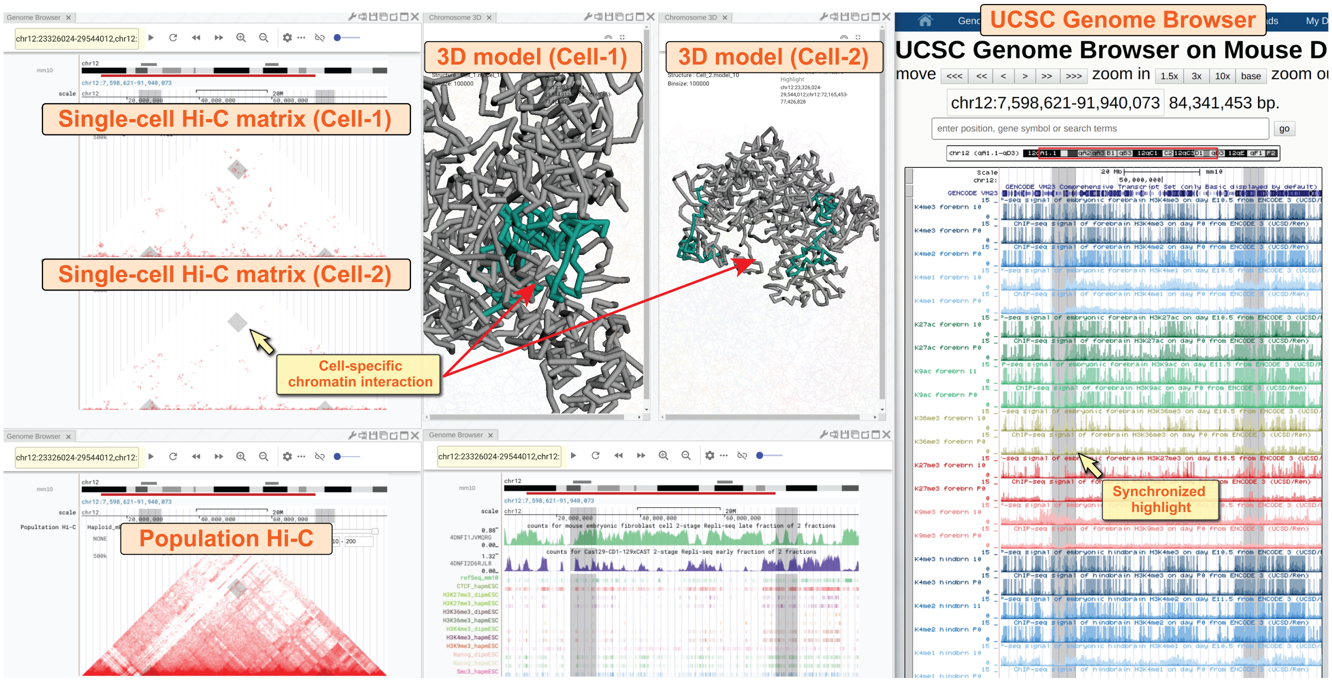


**Figure S22:** An illustration of exploring mouse single-cell Hi-C datasets together with the inferred 3D genome structure models. Two single-cell Hi-C contact maps are shown on the top-left. Hi-C contact map derived from bulk Hi-C is shown on the bottom-left. The 3D genome structures inferred from Hi-C data in these two cells are shown on the center-top, together with ChIP-seq data tracks. The corresponding UCSC Genome Browser view on the same region is shown on the right. Note that the highlighted regions on all the panels including the external UCSC Genome Browser are synchronized, which greatly facilitates multimodal data navigation and comparison.

```
# Medium format (for single cell Hi-C)
<readname> <str1> <chr1> <pos1> <frag1> <str2> <chr2> <pos2> <frag2> <mapq1> <mapq2>

# Short with score format (for population Hi-C, which is already binned)
<str1> <chr1> <pos1> <frag1> <str2> <chr2> <pos2> <frag2> <score>
```

We provide customized script to convert population/single-cell Hi-C contact pairs to .hic files:

```
# Single cell Hi-C
./create_single_cell_HiC.sh Cell_X_contact_pairs.txt Cell_X /path_to_juicer_tools/juicer_tools.jar

# Population Hi-C
./create_population_HiC.sh GSM2123564_Haploid_mESC_population_hic.txt population_hic /path_to_juicer_tools/juicer_tools.jar
```

##### *Pre-processing of 3D genome structures*

The 3D genome structures for each single cell are provided in pdb format, and we will need to convert them into nucle3d format.

```
# pdb to Nucle3d
./pdb2Nucle3d.sh Cell_X_genome_structure_model.pdb Cell_X
```

For each single cell, the pdb file contains 10 structures. Therefore, 10 nucle3d files will be created.

##### *Starting the data service*

We have prepared all data for visualization in Nucleome Browser. Next, we prepare the configuration file to specify the source of data. An example of the configuration Excel file can be downloaded from https://github.com/nucleome/Tutorial-SingeCellHiC/blob/master/single_cell_demo.xlsx. Now we are ready to start a local server using NucleServer.

```
# Start data service using nucleserver
./nucleserver start -i Excel_configuration_file.xlsx
```

Open a web browser and go to: http://vis.nucleome.org, you should see the data in the Nucleome Browser. **Supplementary Fig.** S22 shows a screenshot of visualizing single-cell Hi-C data. In this example, single-cell Hi-C matrices from Cell-1 and Cell-2 together with the population Hi-C contact matrix are shown on the left. The inferred 3D genome structure models for Cell-1 and Cell-2 are shown on the center-top. Other genomic data such as ChIP-seq data is shown on the center-bottom. We highlight a region on the 2D contact map, in which paired reads are only observed in Cell-1 but not in Cell-2. This suggests that the two genomic loci may locate closer to each other in Cell-1 than in Cell-2. In the 3D model, we indeed observe that these two loci are closer to each other in 3D structures in Cell-1 than in Cell-2.

[18] Hubbard T, Barker D, Birney E, Cameron G, Chen Y, Clark L, et al. The Ensembl genome database

project. Nucleic Acids Research. 2002;30(1):38-41.

### C Additional Supplementary Figures and Supplementary Tables

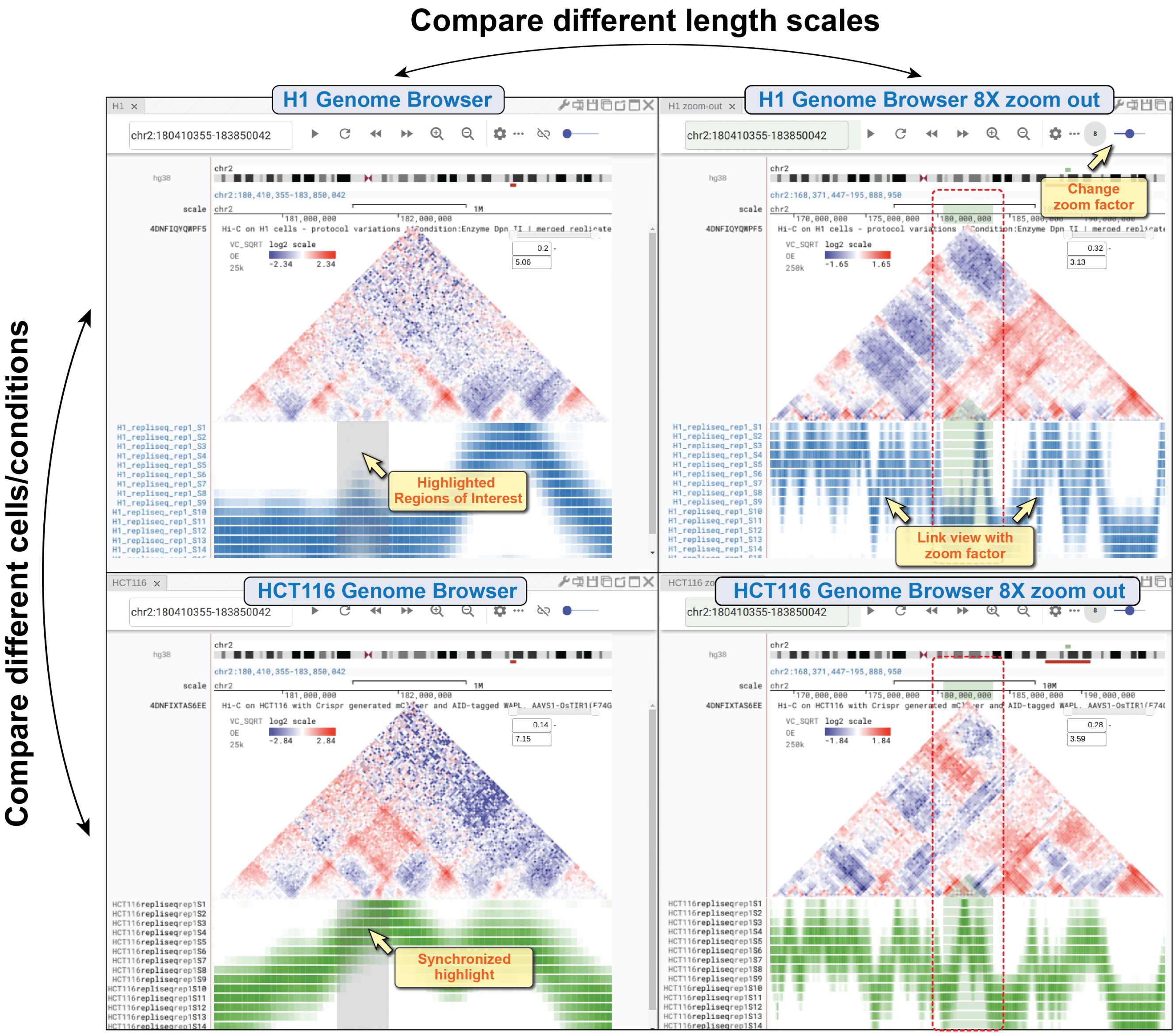


**Figure S23:** Nucleome Browser allows users to create multiple linked panels (e.g., genome browser panels) for interactive exploration. Users can synchronously explore and compare genomic data as shown by the two panels on the left, where an interactive highlighted region indicates an H1 specific early replicated region as compared to HCT116. The two panels on the right are similar to their counterparts on the left except that these panels are in the context mode. In particular, these two panels on the right show the zoomed-out view (8 times) compared to the panels on the left, enabling users to compare genomic data at different scales. Users can use the zoom factor selection tool to easily change zoom factor (from 1x to 64x)

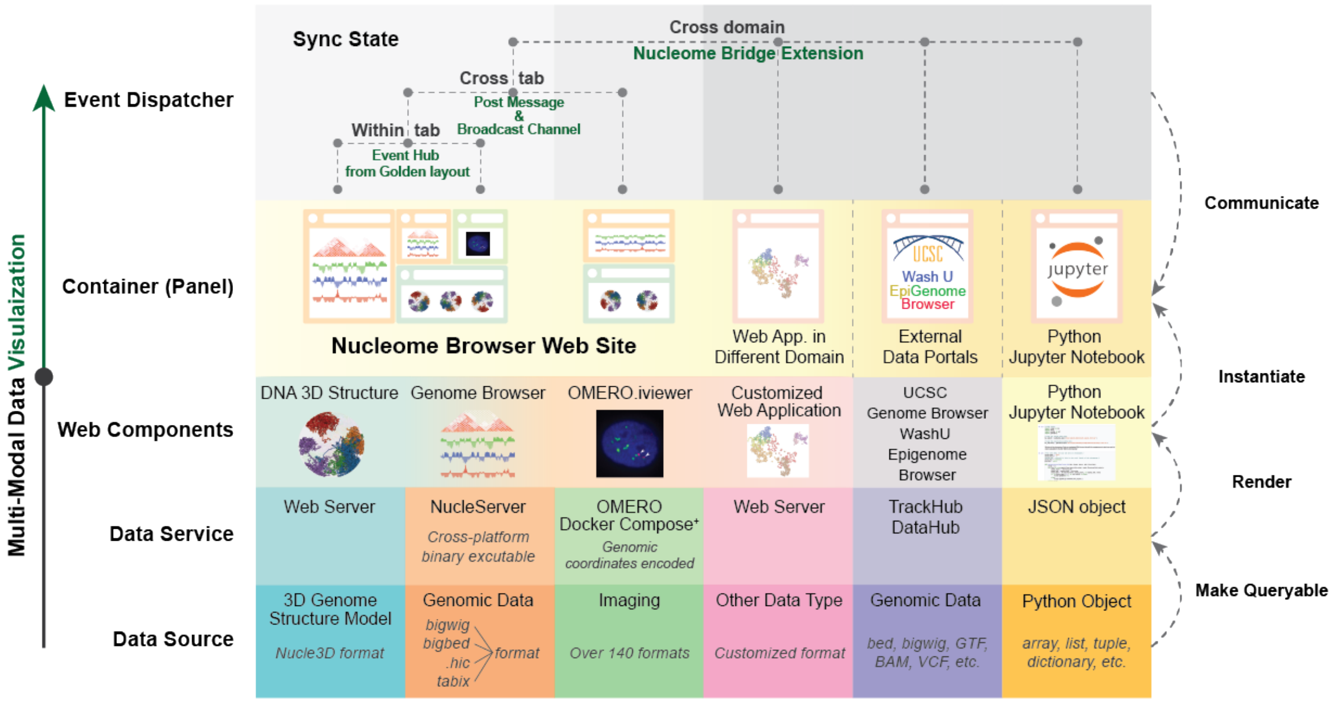


**Figure S24:** The overall design of Nucleome Browser: from data service to synchronization of web components. This diagram shows the key concepts as well as major functionalities introduced in Nucleome Browser to overcome the major challenges of multimodal data visualization for 4D Nucleome research. Each row of the diagram represents a layer of data processing step from bottom to top. Six columns refer to three main modalities (genomics, imaging, and 3D genome structure model) currently represented in Nucleome Browser, customized data type, external data portal, and Python Jupyter Notebook. From bottom to top, it shows the procedure of using data service to process different data types and various front-end to render and visualize these data types. The top two rows are the major components of the adaptive communication system. Web components are represented by composable web panels. We use different web APIs to allow communication between web components in different configurations. Importantly, we also developed Nucleome Bridge web browser extension to communicate with external web-based data portals.

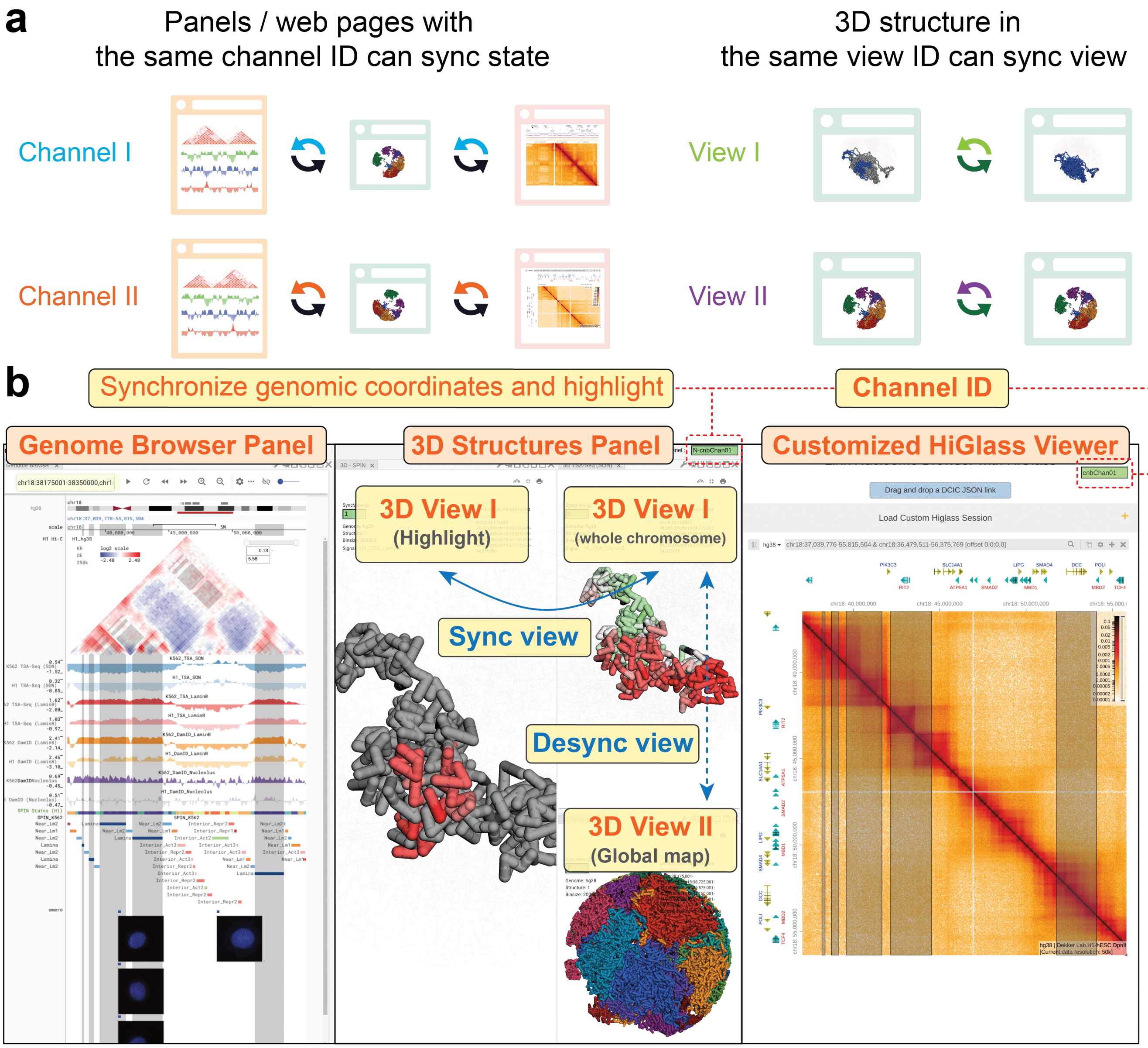


**Figure S25:** Nucleome Browser supports multi-channel communication. **(a)** An illustration of synchronization across panels and 3D genome structure with multi-channel or multi-view communication. Left: panels can only synchronize their operation (e.g., navigation and highlight) to other panels with the same communication channel ID. Right: For the same 3D genome structure, panels with the same view ID can synchronize the rotation and zooming of the 3D structure. **(b)** A screenshot shows the application of the multi-channel and multi-view communication mechanism. The genome browser panel shares the same channel ID with a custom HiGlass viewer page so that the current region and the highlighted regions are synchronized between them. The bottom panel has a view ID different from two 3D genome structure panels on the top. Users can use the bottom panel to view the global reference map of the nucleus and use the other 3D genome structure panel to visualize details such as the highlighted regions and the current chromosome.

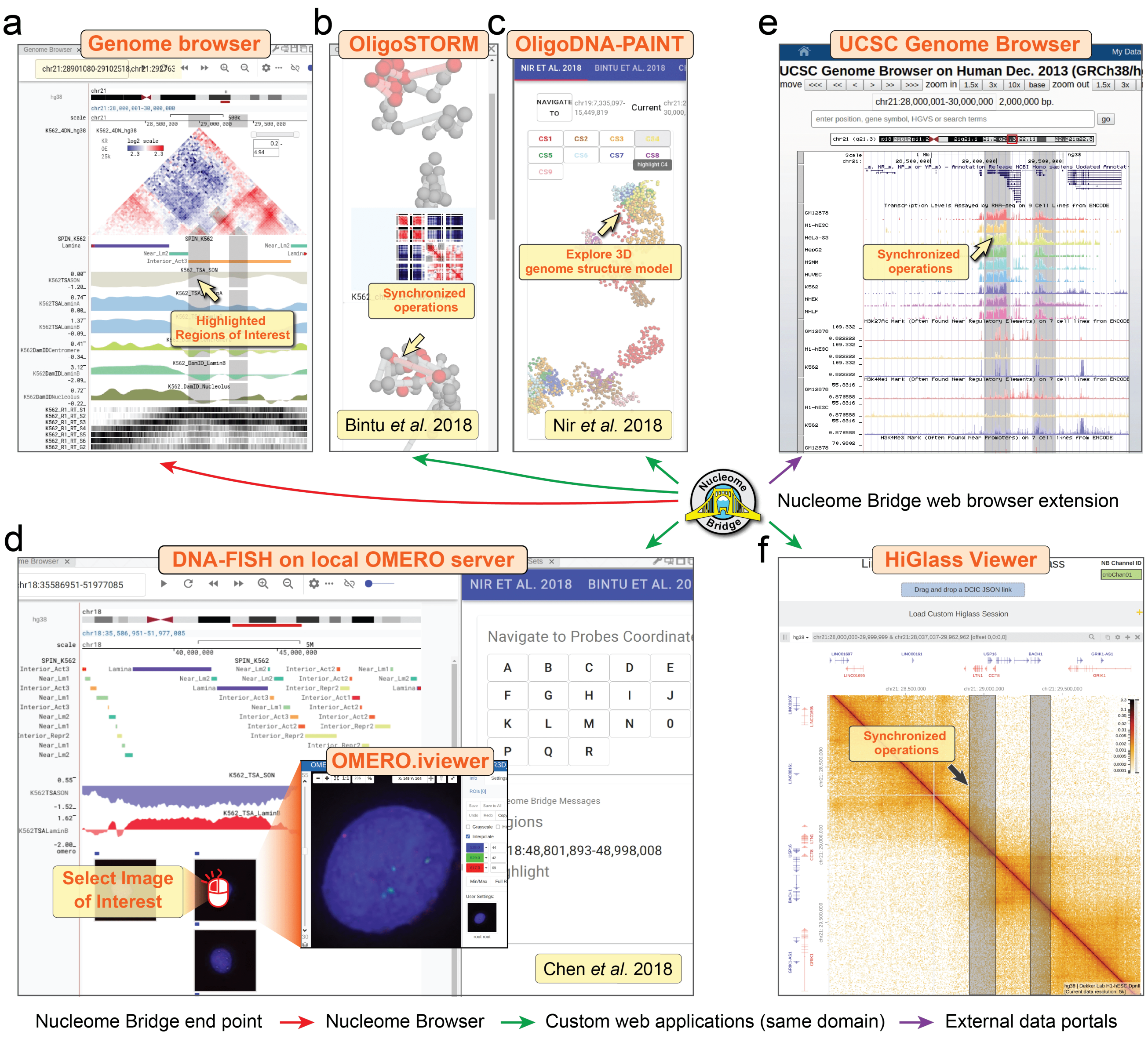


**Figure S26:** Nucleome Browser allows users to build customized web applications to visualize new data types and analysis results together with datasets available in Nucleome Browser. **(a)** When users navigate the genome browser in Nucleome Browser, customized web applications can update their content synchronously. **(b)** This web application shows the 3D genome structure model and 2D distance matrix derived from hundreds of cells using OligoSTORM data (from [7]). Users can select a cell to visualize its 3D model and 2D distance matrix. Probes highlighted in the genome browser in panel a) are marked in red colors in 3D models and highlighted in the 2D distance matrix as well. **(c)** Users can interactively explore 3D genome structure models (spheres indicate probes) derived from OligoDNA-PAINT data (from [8]) and genomic data in Nucleome Browser. Clicking a sphere can navigate to the genomic location of this probe. **(d)** Users can click the button to jump to the probe location in the genome browser on the left. Clicking a thumbnail on the image track will bring up an OMERO.iviewer to show the image data. **(e-f)** Nucleome Browser supports synchronized exploration with external data portals and custom web applications via the new adaptive communication mechanism. When users navigate to or highlight region(s) on the genome browser panel in Nucleome Browser, the connected external data portals (e.g., UCSC Genome Browser, custom web applications, and HiGlass viewer) will update accordingly.

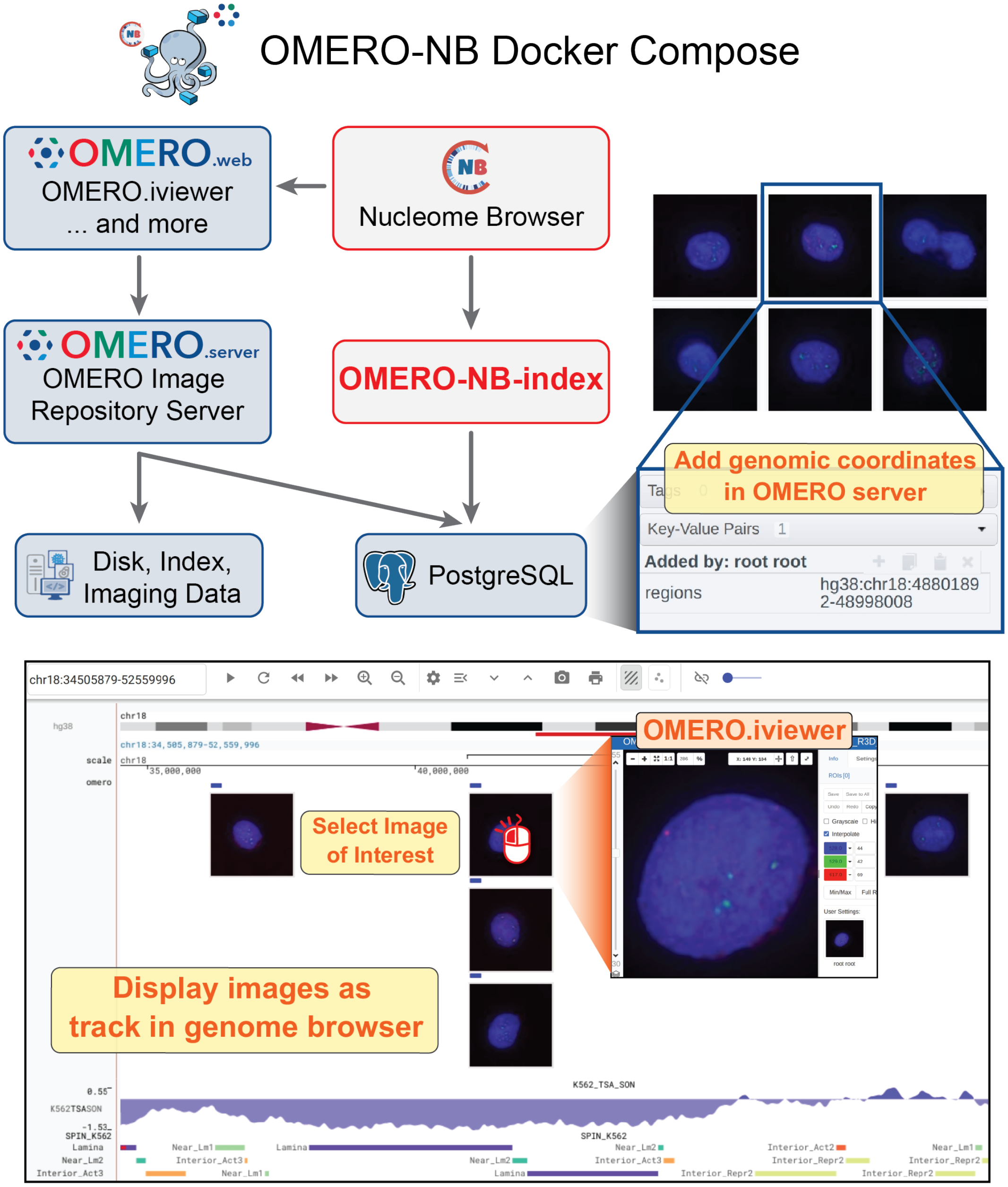


**Figure S27:** Integration between Nucleome Browser and OMERO image server. Top: We built OMERO-NB-index, a specific plugin of OMERO server to link OMERO server with Nucleome Browser. Users can add genomic coordinates associated with images as key:value annotations in the OMERO server. OMERO-NB-index then starts a web server for efficient query of imaging datasets based on genomic coordinates. Bottom: On the genome browser panel in Nucleome Browser, the thumbnails of images are rendered as tracks with red bars indicating the locations of probes. Users then can click a thumbnail to explore the imaging data using the OMERO.iviewer.

| Features | UCSC Genome Browser [1] | Ensembl Genome Browser [18] | WashU Epigenome Browser [3] | IGV & IGV.js [5] | JBrowse [19] | 3D Genome Browser [20] | GIVE [13] | Juicebox & Juicebox.js [21] | HiGlass [6] | Nucleome Browser |
|---|---|---|---|---|---|---|---|---|---|---|
| **Basic functions** | | | | | | | | | | |
| Multiscale visualization | *** | *** | *** | *** | *** | *** | *** | *** | *** | *** |
| Gene annotation | *** | *** | *** | *** | ** | *** | * | ** | *** | *** |
| bigBed/bigWig track | *** | *** | *** | *** | *** | *** | * | *** | *** | *** |
| **4D Nucleome data types** | | | | | | | | | | |
| Intra-chromosomal interaction (Hi-C) | ** | - | * | - | - | ** | * | *** | *** | *** |
| Inter-chromosomal interaction (Hi-C) | - | - | - | - | - | * | * | *** | *** | *** |
| 3D genome structure model | - | - | * | - | - | - | - | - | - | *** |
| **Integration and interactive exploration of imaging data with other data types** | | | | | | | | | | |
| Integrating imaging data from OMERO server | - | - | - | - | - | - | - | - | - | *** |
| Integrating imaging data from Image Data Resource (IDR) | - | - | - | - | - | - | - | - | - | ***, a |
| **Interactive exploration** | | | | | | | | | | |
| Chromosome grid | - | - | - | - | - | - | - | *** | *** | ** |
| Highlight regions on 1D genomic track using mouse | *** | - | - | - | - | - | - | - | - | ***, b |
| Highlight regions on 2D genomic track using mouse | - | - | - | - | - | - | - | - | * | *** |
| Synchronization between windows (navigation/highlight) | - | - | - | - | - | - | - | ** | *** | ***, c |
| Linked windows with variable zoom factors | - | - | - | - | - | - | - | - | * | *** |
| Synchronization between genome browser and 3D structure | - | - | - | - | - | - | - | - | - | *** |
| Super-impose bigWig and bigBed on 3D structure | - | - | - | - | - | - | - | - | - | *** |
| Interactive scatterplot analysis tool | - | - | - | - | - | - | - | - | - | ***, d |
| **Interoperability with other data portal & web services & web applications** | | | | | | | | | | |
| Session URL sharing | *** | *** | *** | * | *** | - | - | *** | *** | *** |
| Set up local service | ** | - | ** | *** | ** | - | *** | *** | *** | *** |
| Connect local service to main server | - | - | - | - | - | - | - | - | - | *** |
| Interactive data analysis with Jupyter Notebook | - | - | - | - | - | - | - | - | - | *** |
| Interactive exploration with external web-based portals | - | - | - | - | - | - | - | - | - | *** |

**Table S1:** Comparing different features between Nucleome Browser with existing genome browsers and 3D genome visualization tools. The support of different tools for each feature is ranked by number of stars from weak to strong. '-' means that this feature is not supported by that tool. **a.** Nucleome Browser can link to IDR to interactively explore data (e.g., the *in situ* genome sequencing (IGS) data) with genome browser. As users highlight regions in the genome browser panels, probes overlapping with highlighted regions are marked on image. WashU Epigenome Browser only supports query image data by gene name based on the description of image datasets in IDR. **b.** Nucleome Browser allows users to highlight genomic regions with the same annotation based on bigBed track, which is not supported on UCSC Genome Browser. **c.** Nucleome Browser allows users to synchronize both navigation and highlight while HiGlass and Juicebox only support navigation. **d.** Nucleome Browser allows users to select data points and highlight corresponding regions on the genome browser. Highlighted regions on the genome browser are also marked in the scatterplot in real time. WashU Epigenome Browser only supports generating static scatterplot without interactive exploration.

| Name | Description | Link |
|---|---|---|
| NucleServer | A Cross-platform tool to launch genomic/3D data service | https://github.com/nucleome/nucleserver |
| NucleData | The GUI version of NucleServer | https://github.com/nucleome/nucledata |
| nb-dispatch | A JavaScript Package of the event-dispatch system | https://github.com/nucleome/nb-dispatch |
| Nucleome Bridge | A Web browser extension for synchronization with UCSC Genome Browser, WashU Epigenome Browser and other customized apps | https://tinyurl.com/nb-bridge |
| OMERO-NB-index | A custom add-on to monitor the image data hosted by the OMERO server and launch a data service for images | https://github.com/nucleome/omero2cnb |
| Nucle3D | The file format used to represent 3D genome structure models in the Nucleome Browser | https://github.com/nucleome/nucle3d |
| NucleTools | The utility scripts for converting data formats for 3D genome structure web component | https://github.com/nucleome/nucle |
| HiGlass viewer page | A customized web page to link HiGlass displays with the Nucleom Browser | https://vis.nucleome.org/static/apps/higlass |
| Nucleome Gallery | The web portal to publish Nucleome Browser sessions | https://gallery.nucleome.org |
| Video tutorial | The ultimate guide on how to use Nucleome Browser | https://tinyurl.com/nb-video-tutorial |
| Documentation | Documentation of Nucleome Browser | https://nb-docs.readthedocs.io |

**Table S2:** Summary of tools/packages/websites created in Nucleome Browser

| Experiment type | Genome | Processed file count |
|---|---|---|
| 2-stage Repli-seq | human | 74 |
| ATAC-seq | human | 18 |
| Capture Hi-C | human | 28 |
| ChIP-seq | human | 24 |
| DamID | human | 44 |
| Dilution Hi-C | human | 267 |
| DNase Hi-C | human | 12 |
| *in situ* Hi-C | human | 766 |
| Micro-C | human | 88 |
| Multi-stage Repli-seq | human | 83 |
| pa-DamID | human | 216 |
| PLAC-seq | human | 11 |
| RNA-seq | human | 84 |
| TSA-seq | human | 44 |
| 2-stage Repli-seq | mouse | 74 |
| ATAC-seq | mouse | 15 |
| Capture Hi-C | mouse | 9 |
| ChIP-seq | mouse | 126 |
| DamID | mouse | 6 |
| Dilution Hi-C | mouse | 28 |
| DNase Hi-C | mouse | 7 |
| *in situ* Hi-C | mouse | 224 |
| Micro-C | mouse | 8 |
| PLAC-seq | mouse | 7 |
| RNA-seq | mouse | 29 |

**Table S3:** Summary of the public 4DN data hosted in Nucleome Browser (synchronized with the 4DN Data Portal (https://data.4dnucleome.org/). Our latest version was updated in November 2021.

| Data set | Data type | Reference or URL |
|---|---|---|
| 4DN public genomic data | Genomic tracks | https://data.4dnucleome.org |
| Single-cell Hi-C | Genomic tracks | [16] |
| 4DN public imaging data | Imaging | https://data.4dnucleome.org |
| *in situ* genome sequencing (IGS) | Imaging | https://idr.openmicroscopy.org/webclient/?show=project-2054, [11] |
| DNA-FISH | Imaging | [22] |
| OligoSTORM | 3D genome structure model | [7] |
| OligoDNA-PAINT | 3D genome structure model | [8] |
| Structure models derived from Hi-C data | 3D genome structure model | [23] |

**Table S4:** Summary of datasets used in this manuscript
